# PCA outperforms popular hidden variable inference methods for molecular QTL mapping

**DOI:** 10.1101/2022.03.09.483661

**Authors:** Heather J. Zhou, Lei Li, Yumei Li, Wei Li, Jingyi Jessica Li

## Abstract

Estimating and accounting for hidden variables is widely practiced as an important step in molecular quantitative trait locus (molecular QTL, henceforth “QTL”) analysis for improving the power of QTL identification. However, few benchmark studies have been performed to evaluate the efficacy of the various methods developed for this purpose. Here we benchmark popular hidden variable inference methods including surrogate variable analysis (SVA), probabilistic estimation of expression residuals (PEER), and hidden covariates with prior (HCP) against principal component analysis (PCA)—a well-established dimension reduction and factor discovery method—via 362 synthetic and 110 real data sets. We show that PCA not only underlies the statistical methodology behind the popular methods but is also orders of magnitude faster, better-performing, and much easier to interpret and use. To help researchers use PCA in their QTL analysis, we provide an R package `PCAForQTL` along with a detailed guide, both of which are freely available at https://github.com/heatherjzhou/PCAForQTL.

## 1 Introduction

Genome-wide association studies (GWASs) have identified thousands of genetic variants associated with human traits or diseases [1–4]. However, the majority of GWAS variants are located in non-coding regions of the genome, making it challenging to interpret the GWAS associations [5, 6]. In response to this, molecular quantitative trait locus (molecular QTL, henceforth “QTL”) analysis has emerged as an important field in human genetics, interrogating the relationship between genetic variants and intermediate, molecular traits and potentially explaining the GWAS associations [7, 8].

Based on the type of molecular phenotype studied, QTL analyses can be categorized into gene expression QTL (eQTL) analyses [9, 10], alternative splicing QTL (sQTL) analyses [10], three prime untranslated region alternative polyadenylation QTL (3^′^aQTL) analyses [11], and so on [7, 8]. Among these categories, eQTL analyses, which investigate the association between genetic variants and gene expression levels, are the most common. To date, most (single-tissue) QTL studies are carried out using regression-based methods such as Matrix eQTL [12] and FastQTL [13].

In QTL analysis, a major challenge is that measurements of gene expression levels and other molecular phenotypes can be affected by a number of technical or biological variables other than the genetic variants (e.g., batch, sex, and age). If these variables are known, then they can be directly included in the QTL pipeline as covariates. However, many of these variables may be unknown or unmeasured. Therefore, it has become standard practice to *first* infer the hidden variables and *then* include the inferred variables as covariates or otherwise account for them (Table 1; see Section 4.3 for a numerical example) [9–11, 14–23]. This type of approach has been shown to both improve the power of QTL identification in simulation settings [24] and empirically increase the number of discoveries in QTL studies [9, 10, 16, 21–23].

**Table 1:** Summary of the 15 methods we compare based on simulation studies, including Ideal, Unadjusted, and 13 variants of PCA, SVA, PEER, and HCP (Section S4). Out of the 15 methods, we select a few representative methods (Section 4.2) for detailed comparison in Simulation Design 2, the abbreviations of which are shown in (D). *Y* denotes the gene expression matrix, *Y*_resid_ denotes the residual matrix outputted by PEER, *X*_1_ denotes the known covariate matrix, and *X*_2_ denotes the hidden covariate matrix. In Line 3, PCA is run on *Y* directly; in Line 4, PCA is run after the effects of *X*_1_ are regressed out from *Y* (Section S4). The addition signs in (C) denote column concatenation. “filtered” means that we filter out the known covariates that are captured well by the inferred covariates (unadjusted *R*^2^ ≥ 0.9); this filtering is only needed when the hidden variable inference method in (A) does not explicitly take the known covariates into account.

|  | Inference method | Method | Response, covariates | Method abbr. (if selected) |
| --- | --- | --- | --- | --- |
|  | (A) | (B) | (C) | (D) |
| 1 | | **Ideal** | $Y, X_1 + X_2$ | Ideal |
| 2 | | **Unadjusted** | $Y, X_1$ | Unadjusted |
| 3 | PCA_direct | **PCA_direct_sceeK** | $Y, X_1$ (filtered) + top PCs | PCA |
| 4 | PCA_resid | **PCA_resid_sceeK** | $Y, X_1$ + top PCs | |
| 5 | SVA_trueK | **SVA_trueK** | $Y, X_1$ + SVs | |
| 6 | SVA_BE | **SVA_BE** | $Y, X_1$ + SVs | SVA |
| 7 | PEER_noCov_trueK | **PEER_noCov_trueK_factors** | $Y, X_1$ (filtered) + PEER factors | |
| 8 | | **PEER_noCov_trueK_residuals** | $Y_{\text{resid}}, \text{NULL}$ | |
| 9 | PEER_noCov_largeK | **PEER_noCov_largeK_factors** | $Y, X_1$ (filtered) + PEER factors | |
| 10 | | **PEER_noCov_largeK_residuals** | $Y_{\text{resid}}, \text{NULL}$ | |
| 11 | PEER_withCov_trueK | **PEER_withCov_trueK_factors** | $Y, X_1$ + PEER factors | PEER, true K, factors |
| 12 | | **PEER_withCov_trueK_residuals** | $Y_{\text{resid}}, \text{NULL}$ | |
| 13 | PEER_withCov_largeK | **PEER_withCov_largeK_factors** | $Y, X_1$ + PEER factors | |
| 14 | | **PEER_withCov_largeK_residuals** | $Y_{\text{resid}}, \text{NULL}$ | PEER, large K, residuals |
| 15 | HCP_trueK | **HCP_trueK** | $Y, X_1$ + HCPs | HCP |

Surrogate variable analysis (SVA) [25, 26] is one of the first popular hidden variable inference methods for large-scale genomic analysis. Although initially proposed as a hidden variable inference method for both QTL mapping and differential expression (DE) analysis, currently SVA is primarily used in DE and similar analyses as opposed to QTL mapping [27–30]. We believe this is partly because the SVA package [31] is difficult to apply in QTL settings in that it requires the user to input at least one variable of interest and using too many variables of interest causes the package to fail (Figure 1; Section S4); while there are usually at most a few variables of interest in a DE study, there are often millions of single nucleotide polymorphisms (SNPs; variables of interest) in a QTL study. Historically, there have been two versions of the SVA method: two-step SVA [25] and iteratively reweighted SVA (IRW-SVA) [26]; the latter supersedes the former. Therefore, we focus on IRW-SVA in this work.

**Figure 1:**
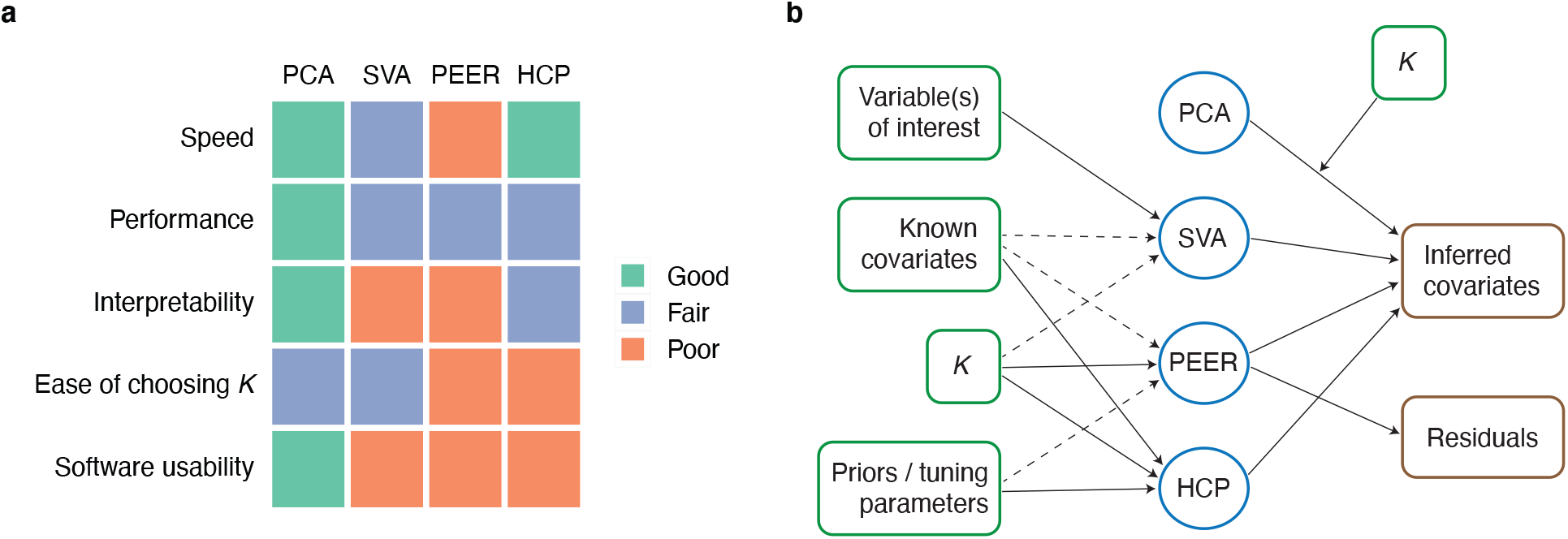
Overall comparison of PCA, SVA, PEER, and HCP and summary of their inputs and outputs. In this work, we use *K* to denote the number of inferred covariates, which are called PCs, SVs, PEER factors, and HCPs in PCA, SVA, PEER, and HCP, respectively. **a** PCA is faster, better-performing, and much easier to interpret and use. For speed and performance comparison, see Section 2.1 (and to a lesser extent, Sections 2.2 and 2.3). For interpretability and ease of choosing *K*, see Sections 2.4 and 2.5, respectively. In terms of software usability, SVA is difficult to apply in QTL settings (Section S4), PEER is difficult to install, and HCP is poorly documented. In addition, PEER suffers from the disadvantage that there is no consensus in the literature on how it should be used (Section S4). **b** Inputs (green boxes) and outputs (brown boxes) of the four methods. The fully processed molecular phenotype matrix (after the effects of the known covariates are regressed out in the case of PCA resid; Table 1) is a required input for all four methods and is thus omitted in the diagram. Dashed arrows indicate optional inputs. PEER outputs both inferred covariates and residuals of the inputted molecular phenotype matrix [32].

Probabilistic estimation of expression residuals (PEER) [24, 32] is currently the most popular hidden variable inference method for QTL mapping by far. It is used in the Genotype-Tissue Expression (GTEx) project [9, 10] and many other high-impact studies [11, 14–21]. The PEER method has two main perceived advantages: (1) it can take known covariates into account when estimating the hidden covariates, and (2) its performance does not deteriorate as the number of inferred covariates increases (i.e., it does not “overfit”). One drawback of PEER, though, is that there is no consensus in the literature on *how* it should be used. For example, when there are known covariates available, PEER can be run with or without the known covariates—Stegle et al. [32] do not give an explicit recommendation as to which approach should be used, and both approaches are used in practice (e.g., [9, 10] vs. [11, 16]). Further, PEER outputs both inferred covariates and residuals of the inputted molecular phenotypes (Figure 1), so the user needs to decide which set of outputs to use (Section S4; we refer to the approach using the inferred covariates as the “factor approach” and the approach using the residuals as the “residual approach”). Such “flexibility” of PEER could be considered a benefit, but we believe it not only leads to confusion for practitioners but also reduces the transparency and reproducibility of published QTL research.

Hidden covariates with prior (HCP) [33] is another popular hidden variable inference method for QTL mapping. Though less popular than PEER, it is also used in some high-impact studies [22, 23]. To determine which method is the best and whether PEER indeed has the perceived advantages, we thoroughly evaluate SVA, PEER, and HCP for the first time in the literature. Given that principal component analysis (PCA) [34–38] underlies the methodology behind each of these methods (Section 2.4) and has indeed been used for the same purpose [39, 40], we also include PCA in our evaluation. Through simulation studies (Section 2.1) and real data analysis (Sections 2.2, 2.3 and 2.5), we show that PCA is orders of magnitude faster, better-performing, and much easier to interpret and use (Figure 1).

## 2 Results

### 2.1 Comprehensive simulation studies show that PCA is faster and better-performing

We compare the runtime and performance of 15 methods (Table 1), including Ideal (assuming the hidden covariates are known), Unadjusted (not estimating or accounting for the hidden covariates), and 13 variants of PCA, SVA, PEER, and HCP, based on two simulation studies. In the first simulation study (Simulation Design 1; Section S2), we follow the data simulation in Stegle et al. [24]—the original PEER publication—while addressing its data analysis and overall design limitations (Section S1). In the second simulation study (Simulation Design 2; Section S3), we further address the data simulation limitations of Stegle et al. [24] (Section S1) by simulating the data in a more realistic and comprehensive way, roughly following Wang et al. [41]—the SuSiE publication—but introducing the existence of known and hidden covariates. A summary of the main differences between the two simulation designs is provided in Table S1. The key difference is that in Simulation Design 1, the gene expression levels are primarily driven by trans-regulatory effects rather than cis-regulatory effects or covariate effects (Table S2), inconsistent with the common belief that trans-regulatory effects are generally weaker than cis-regulatory effects. In contrast, in Simulation Design 2, we focus on cis-QTL detection and carefully control the genotype effects and covariate effects in 176 experiments with two replicates per experiment (Section S3).

The details of the 15 methods are described in Section S4, and the evaluation metrics are described in Section 4.1. For convenience, we refer to the simulated molecular phenotypes as gene expression levels throughout our simulation studies; however, they can be interpreted as any type of molecular phenotype after data preprocessing and transformation, e.g., alternative splicing phenotypes and alternative polyadenylation phenotypes (Table S3).

The results from our simulation studies are summarized in Figures 2, 3, S1, S3, and S4. We find that PCA and HCP are orders of magnitude faster than SVA, which in turn is orders of magnitude faster than PEER, and that PCA outperforms SVA, PEER, and HCP in terms of the area under the precision-recall curve (AUPRC) of the QTL result (Figures 2 and 3). On a dataset-by-dataset basis, PCA outperforms the other methods in terms of AUPRC in 11% to 88% of the simulated data sets and underperforms them in close to 0% of the simulated data sets in Simulation Design 2 (Figure S3(d)). In addition, PCA has the highest average concordance score(s), a metric for the concordance between the true hidden covariates and the inferred covariates (Section 4.1; Figures S1 and S4), which explains why PCA performs the best in terms of AUPRC.

**Figure 2:**
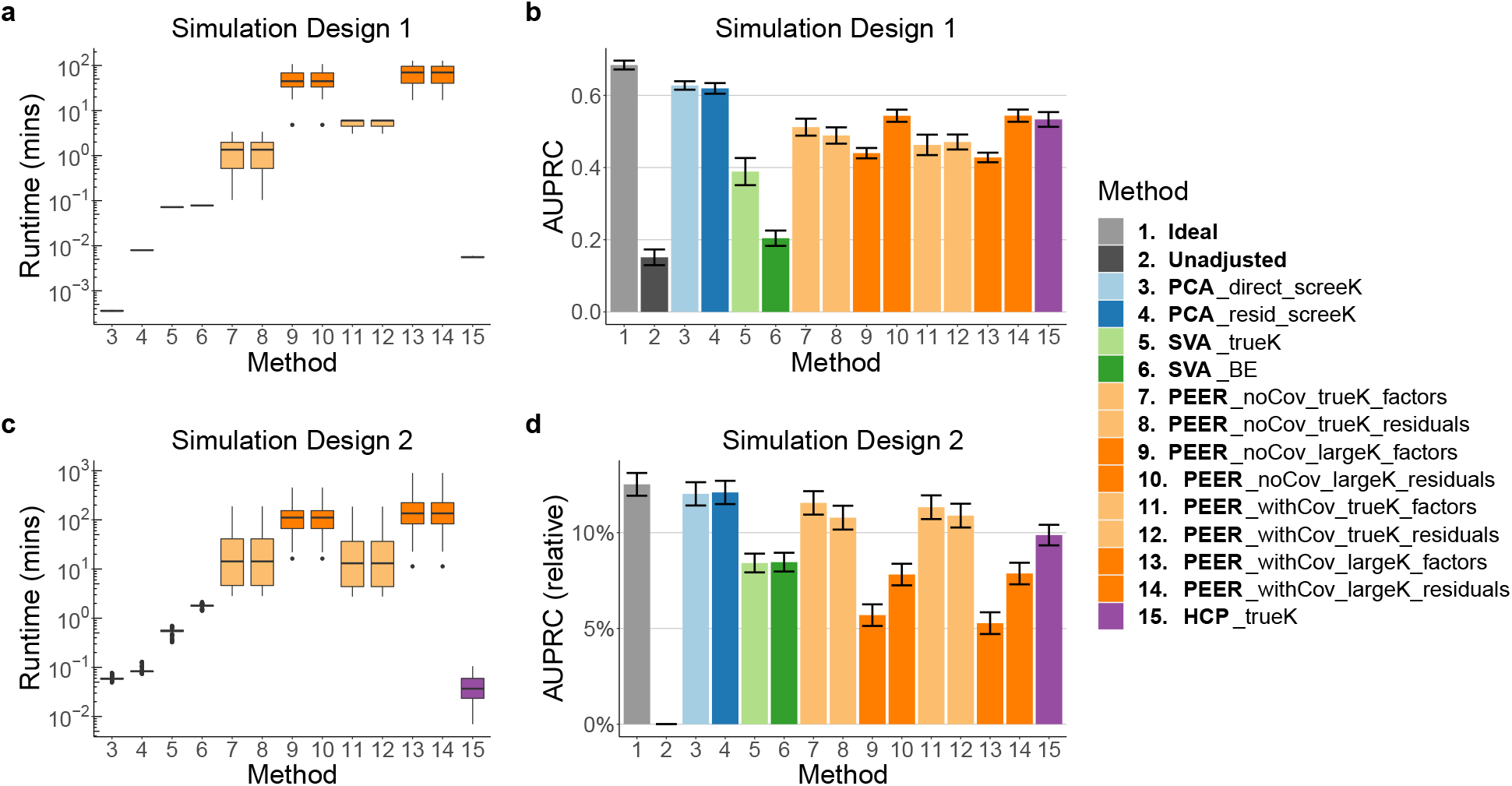
Runtime and AUPRC comparison of all 15 methods (Table 1) in Simulation Design 1 and Simulation Design 2. **a, c** PCA and HCP each takes within a few seconds, SVA takes up to a few minutes, and PEER takes up to about 1, 000 minutes, equivalent to about 17 hours. In particular, PEER takes longer to run when *K* is larger (dark orange vs. light orange boxes). **b, d** PCA outperforms SVA, PEER, and HCP in terms of AUPRC. The height of each bar represents the average across simulated data sets. For ease of visualization, in **d**, the *y*-axis displays AUPRC −AUPRC_Unadjusted_ */*AUPRC_Unadjusted_. In this work, error bars indicate standard errors unless otherwise specified (whiskers in box plots are not considered error bars).

**Figure 3:**
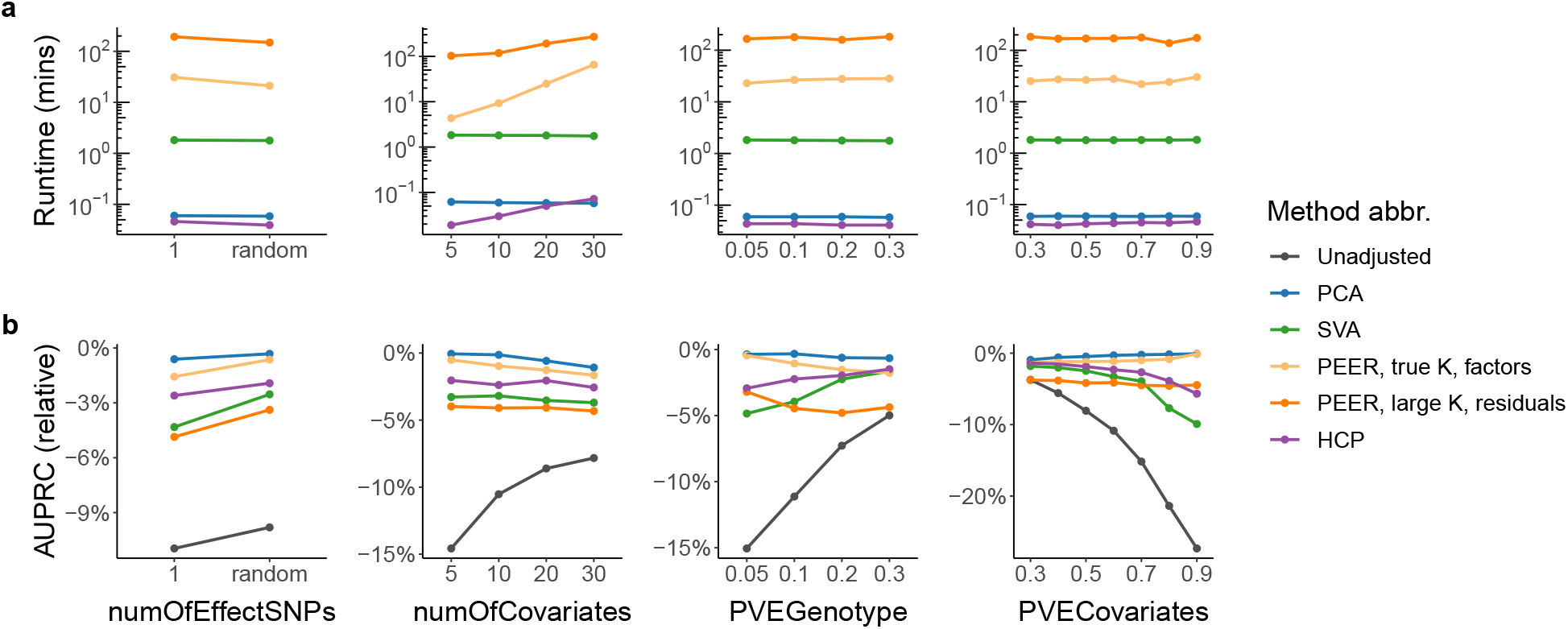
Detailed runtime and AUPRC comparison of the selected representative methods (Table 1) in Simulation Design 2. Each point represents the average across simulated data sets. The *x*-axes are: number of effect SNPs per gene (`numOfEffectSNPs`), number of simulated covariates (`numOfCovariates`; including known and hidden covariates), proportion of variance explained by genotype (`PVEGenotype`), and proportion of variance explained by covariates (`PVECovariates`) (Section S3). **a** PCA and HCP are orders of magnitude faster than SVA, which in turn is orders of magnitude faster than PEER. **b** PCA outperforms SVA, PEER, and HCP in terms of AUPRC across different simulation settings. For ease of visualization, the *y*-axis displays (AUPRC −AUPRC_Ideal_) */*AUPRC_Ideal_. Consistent with our expectation, the performance gap between Unadjusted and Ideal is the largest (and thus accounting for hidden covariates is the most important) when `numOfCovariates` is small, when `PVEGenotype` is small, and when `PVECovariates` is large.

To contrast the results in Stegle et al. [24], we also compare the powers of the different methods in Simulation Design 1 (Figure S1). We find that PCA is more powerful than SVA, PEER, and HCP. Notably, SVA and PEER have very low power in identifying trans-QTL relations—an especially unfavorable result for SVA and PEER, considering that the gene expression levels are primarily driven by trans-regulatory effects in Simulation Design 1 (Table S2).

Incidentally, Figures 2 and S3 also provide us with the following insights into the different ways of using PEER (Section S4). First, running PEER *with* the known covariates has no advantage over running PEER *without* the known covariates in terms of AUPRC, given the choice of *K* (the number of inferred covariates) and the choice between the factor approach and the residual approach. In fact, running PEER *with* the known covariates significantly increases the runtime of PEER in real data (Section 2.3). Second, contrary to claims in Stegle et al. [24, 32], the performance of PEER *does* deteriorate as the number of PEER factors increases. The only exception is when the residual approach is used in Simulation Design 1 (Figure 2). But given that Simulation Design 2 is more realistic than Simulation Design 1 and that the factor approach is more popular than the residual approach [9–11, 17–20], the take-home message should be that in general, the performance of PEER is worse when we use a large *K* rather than the true *K*. Third, whether the factor approach or the residual approach performs better depends on the choice of *K*. When we use the true *K*, the factor approach performs better, but when we use a large *K*, the residual approach performs better. All in all, PCA outperforms all different ways of using PEER in both of our simulation studies (Figure 2).

### 2.2 PEER factors sometimes fail to capture important variance components of the molecular phenotype data

For our real data analysis, we examine the most recent GTEx eQTL and sQTL data [10] (Sections 2.3 and 2.5) and the 3^′^aQTL data prepared by Li et al. [11] from GTEx RNA-seq reads [9] (Section 2.2). While the exact data analysis pipelines are different (Table S3), these studies all choose PEER as their hidden variable inference method.

Unlike PCs, which are always uncorrelated (Section S5.1), PEER factors are not guaranteed to be uncorrelated. Here we show through the above-mentioned 3^′^aQTL data that PEER factors can be highly correlated with each other (to the extent that many or all of them are practically identical) and thus fail to capture important variance components of the molecular phenotype data.

Given a post-imputation alternative polyadenylation phenotype matrix (each entry is between zero and one, representing a proportion), Li et al. [11] run PEER without further data transformation using the number of PEER factors chosen by GTEx [9] (Table S3). To assess the impact of data transformation on the PEER factors, we also run PEER after transforming the data in three ways: (1) center and scale (to unit variance) each feature, (2) apply inverse normal transform (INT) [42] to each feature (“INT within feature”), and (3) apply INT to each sample (“INT within sample”). Among these methods, GTEx [9, 10] uses “INT within feature” for its eQTL data and “INT within sample” for its sQTL data (Table S3). To quantify how many “distinct” or “nonrepetitive” PEER factors there are, given a set of PEER factors, we group them into clusters such that in each cluster, the correlation between any two PEER factors is above a pre-defined threshold (0.99, 0.9, or 0.8) in absolute value (this is done via hierarchical clustering [43] with complete linkage and the distance defined as one minus the absolute value of the correlation). Therefore, the number of PEER factor clusters can be interpreted as the number of distinct or nonrepetitive PEER factors.

Our results show that in many cases, the number of distinct PEER factors is considerably smaller than the number of PEER factors requested (Figure 4), and when this issue is severe (e.g., “No transformation” and “INT within sample”), the PEER factors fail to capture important variance components of the molecular phenotype data (Figure S5). Since the number of discoveries increases substantially with the number of PEER factors in GTEx’s eQTL analyses [9, 10], where the PEER factors are essentially identical to PCs (Section 2.3), it is possible that replacing the nearly-all-identical PEER factors with appropriate numbers of PCs in Li et al. [11]’s 3^′^aQTL analysis can lead to more discoveries. This is a potential direction for a future study.

**Figure 4:**
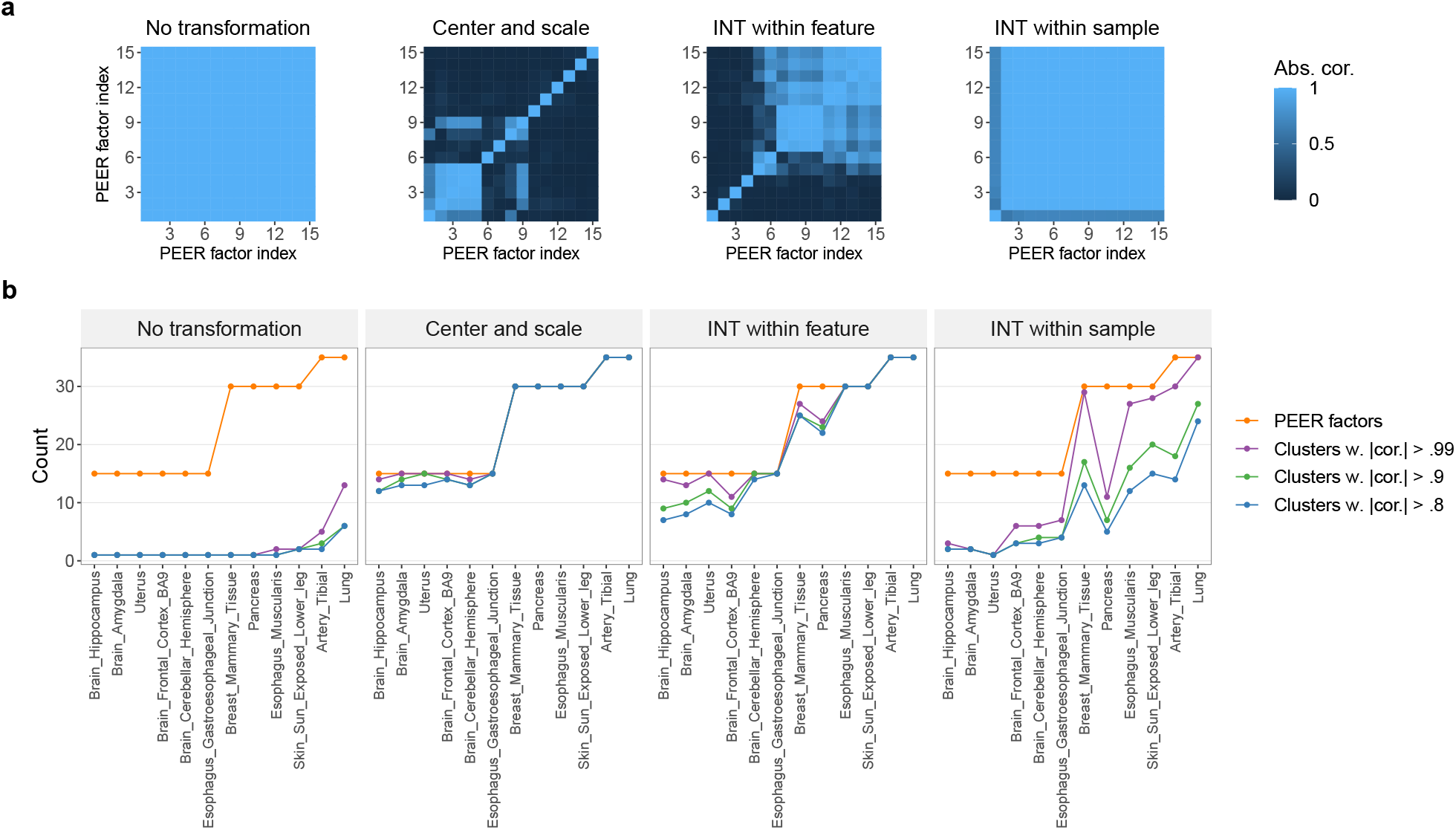
In the 3^′^aQTL data prepared by Li et al. [11] from GTEx RNA-seq reads [9], PEER factors can be highly correlated with each other to the extent that many or all of them are practically identical. **a** Correlation heatmaps of PEER factors for Brain Hippocampus. For ease of visualization, the PEER factors are reordered based on results from hierarchical clustering (Section 2.2). **b** The *x*-axis shows 12 randomly selected tissue types with increasing sample sizes. The *y*-axis shows the number of PEER factors requested (orange line) or the number of PEER factor clusters. In each cluster, the correlation between any two PEER factors is above 0.99, 0.9, or 0.8 in absolute value. Therefore, the number of PEER factor clusters can be interpreted as the number of distinct or nonrepetitive PEER factors. We find that in many cases, the number of distinct PEER factors is considerably smaller than the number of PEER factors requested, and when this issue is severe (e.g., “No transformation” and “INT within sample”), the PEER factors fail to capture important variance components of the molecular phenotype data (Figure S5).

### 2.3 PEER factors are almost identical to PCs but take three orders of magnitude longer to compute in GTEx eQTL and sQTL data

We report the surprising finding that in both GTEx eQTL and sQTL data [10], the PEER factors obtained by GTEx and used in its QTL analyses are almost identical to PCs. Specifically, given a fully processed molecular phenotype matrix, there is almost always a near-perfect one-to-one correspondence between the PEER factors and the top PCs (Figure 5). This means that after the variational Bayesian inference in PEER initializes with PCs [24], it does not update the PCs much beyond scaling them (see Section 2.4 for an explanation). Therefore, it is no surprise that replacing the PEER factors with PCs in GTEx’s FastQTL pipeline [10, 13] does not change the QTL results much (Figures S6 and S7) because in linear regressions (the basis of both Matrix eQTL [12] and FastQTL [13]), scaling and/or shifting the predictors does not change the *p*-values of *t*-tests for non-intercept terms (neither does scaling and/or shifting the response, for that matter).

**Figure 5:**
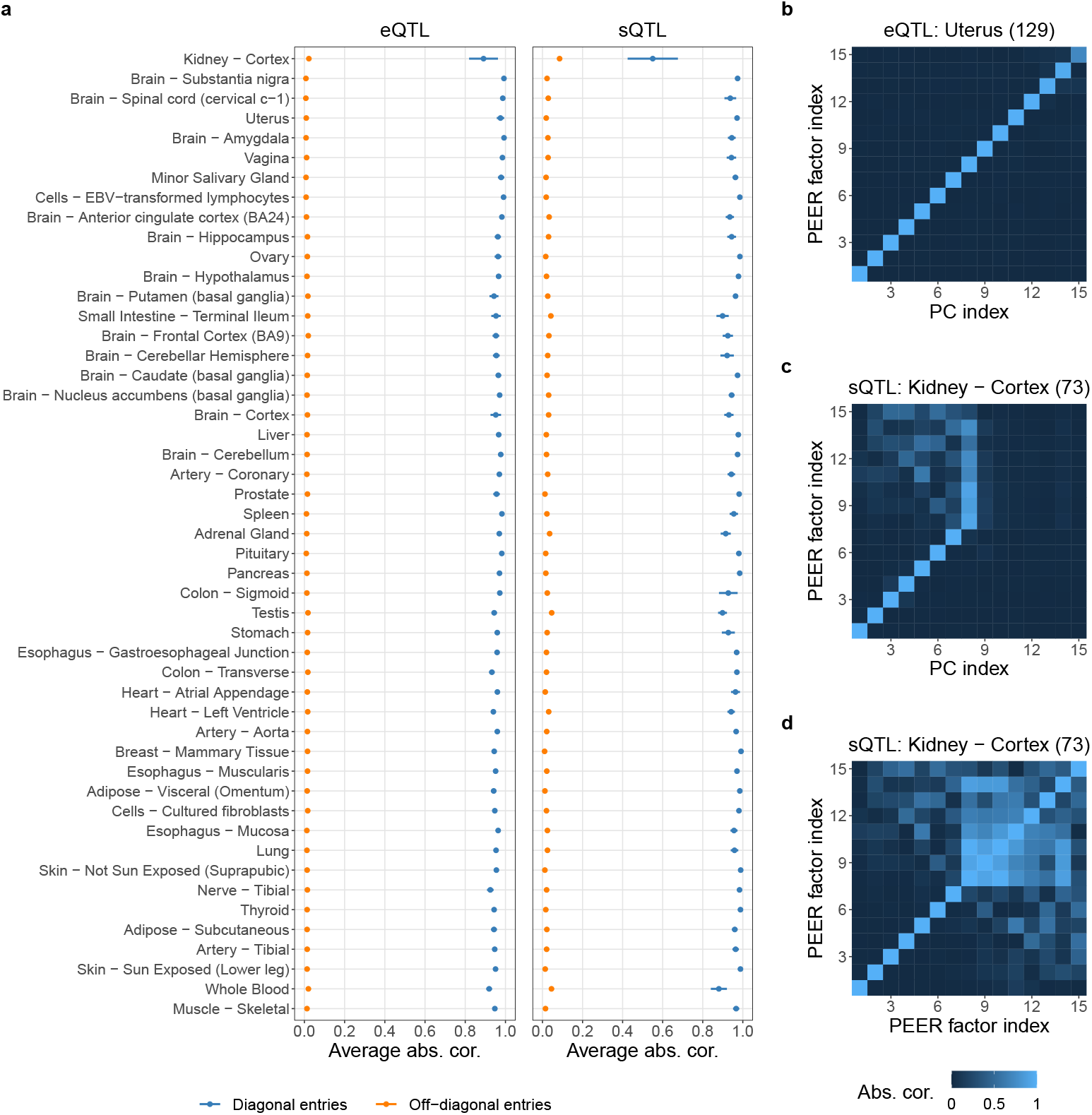
PEER factors are almost identical to PCs in GTEx eQTL and sQTL data [10]. **a** The *y*-axis shows all 49 tissue types with GTEx QTL analyses ordered by sample size (from small to large). Given a fully processed molecular phenotype matrix, we summarize the correlation matrix (in absolute value) between the PEER factors and the top PCs into two numbers: the average of the diagonal entries and the average of the off-diagonal entries. With the exception of Kidney - Cortex sQTL data, the diagonal entries have averages close to one, and the off-diagonal entries have averages close to zero (both have minimal standard errors). **b** A typical correlation heatmap showing near-perfect one-to-one correspondence between the PEER factors and the top PCs. **c** In Kidney - Cortex sQTL data, the PEER factors and the top PCs do not have a perfect one-to-one correspondence. The reason is because the PEER factors are highly correlated with each other (**d**), while PCs are always uncorrelated (Section S5.1). The numbers in the parentheses represent sample sizes. To produce this figure, we reorder the PEER factors based on the PCs (Algorithm S1), although in almost all cases, this reordering does not change the original ordering of the PEER factors because PEER initializes with PCs [24].

However, PEER is at least three orders of magnitude slower than PCA (Figure S6). For a given expression matrix, running PEER without the known covariates (GTEx’s approach) takes up to about 32 hours, while running PCA (with centering and scaling; our approach) takes no more than a minute.

To draw a connection between the simulation results and real data results, we analyze them jointly in Figure S8 and make the following two key observations. First, we find that in the simulation studies, PCA almost always outperforms PEER in terms of AUPRC (confirming our results in Section 2.1), and the percentage of QTL discoveries shared between PEER and PCA is a good predictor of the relative performance of PEER versus PCA—the higher the percentage of QTL discoveries shared, the smaller the performance gap between PEER and PCA. Second, the percentages of QTL discoveries shared between the two methods in GTEx eQTL data [10] fall comfortably within the range of percentage of QTL discoveries shared in Simulation Design 2. These two observations together suggest that PCA likely outperforms PEER in GTEx eQTL data [10] even though the results largely overlap.

### 2.4 PCA, SVA, PEER, and HCP are closely related statistical methods

We report that PCA, SVA, PEER, and HCP are closely related statistical methods despite their apparent dissimilarities. In particular, the methodology behind SVA, PEER, and HCP can all be traced back to PCA (Figure 6). We have previously reviewed these methods in detail in Zhou [44]. Here we aim to provide a brief summary and highlight their connections.

**Figure 6:**
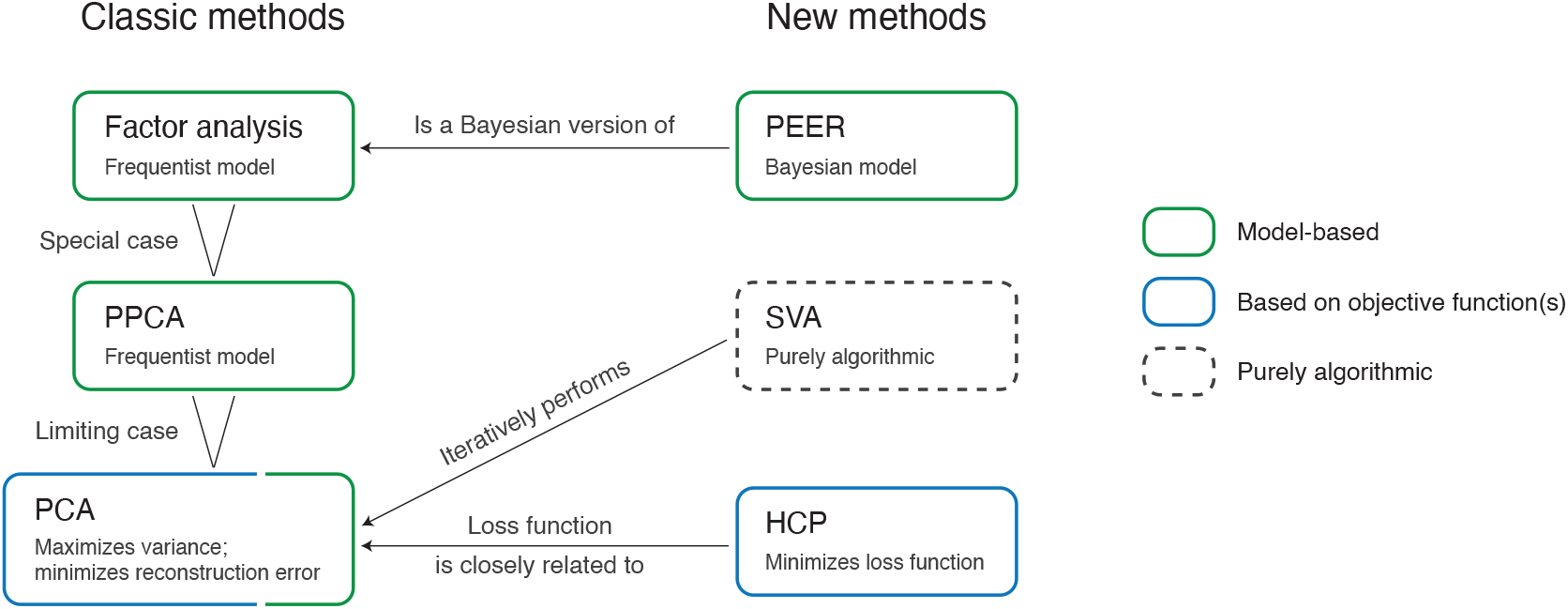
PCA, SVA, PEER, and HCP are closely related statistical methods despite their apparent dissimilarities. In particular, the methodology behind SVA, PEER, and HCP can all be traced back to PCA. PCA [34–38] is traditionally derived by optimizing some objective functions (either maximum variance or minimum reconstruction error; Section S5.1), but more recently, it is shown that PCA can be derived as a limiting case of probabilistic principal component analysis (PPCA) [45], which in turn is a special case of factor analysis [35, 46]. PEER [24, 32] is based on a Bayesian probabilistic model and can be considered a Bayesian version of factor analysis. SVA [25, 26] is purely algorithmic and is not defined based on a probabilistic model or objective function. The steps of the SVA algorithm are complicated [44], but in a nutshell, SVA iterates between two steps: (1) reweight the features of the molecular phenotype matrix, and (2) perform PCA on the resulting matrix (with centering but without scaling) [26]. Lastly, HCP [33] is defined by minimizing a loss function that is very similar to the minimum-reconstruction-error loss function of PCA (Section S5.2).

PCA [34–38] is traditionally derived by optimizing some objective functions (either maximum variance or minimum reconstruction error; Section S5.1), but more recently, it is shown that PCA can be derived as a limiting case of probabilistic principal component analysis (PPCA) [45], which in turn is a special case of factor analysis [35, 46]—a dimension reduction method commonly used in psychology and the social sciences that is based on a *frequentist* probabilistic model.

PEER [24, 32] is based on a *Bayesian* probabilistic model and can be considered a Bayesian version of factor analysis (with the not-very-useful ability to explicitly model the known covariates; see Section 2.1 for why we do not find this ability useful). Inference is performed using variational Bayes and initialized with the PCA solution [24]. Given that PCA underlies the PEER model (Figure 6) and PEER initializes with PCs, it is not surprising that PEER factors are almost identical to PCs in GTEx eQTL and sQTL data [10] (Section 2.3).

SVA [25, 26] is purely algorithmic and is not defined based on a probabilistic model or objective function. The steps of the SVA algorithm are complicated [44], but in a nutshell, SVA iterates between two steps: (1) reweight the features of the molecular phenotype matrix, and (2) perform PCA on the resulting matrix (with centering but without scaling) [26].

Lastly, HCP [33] is defined by minimizing a loss function that is very similar to the minimum-reconstruction-error loss function of PCA (Section S5.2). The optimization is done through coordinate descent with one deterministic initialization (see source code of the HCP R package [33]). In short, SVA, PEER, and HCP can all be considered extensions or more complex versions of PCA, though we show that the complexity is a burden rather than a benefit (Figure 1).

### 2.5 PCA provides insight into the choice of *K*

Choosing *K*, the number of inferred covariates in the context of hidden variable inference or the number of dimensions or clusters in more general contexts, is always a difficult task. Nonetheless, based on the proportion of variance explained (PVE) by each PC (Section S5.1), PCA offers convenient ways of choosing *K* such as the elbow method and the Buja and Eyuboglu (BE) algorithm [47] (more details below). Since SVA is heavily based on PCA (Section 2.4), it is able to adapt and make use of the BE algorithm. In contrast, PEER and HCP do not offer easy ways of choosing *K*; for lack of a better method, users of PEER and HCP often choose *K* by maximizing the number of discoveries [9, 10, 16, 21–23]. Not only is this approach of choosing *K* extremely computationally expensive and theoretically questionable, here we also show from the perspective of PCA that it may yield inappropriate choices of *K*.

Recall from Section 2.3 that PEER factors are almost identical to PCs in GTEx eQTL data [10] (the number of PEER factors is chosen by maximizing the number of discovered cis-eGenes for each pre-defined sample size bin; Table S3). Therefore, for each tissue type, we compare the number of PEER factors selected by GTEx to (1) the number of PCs chosen via an automatic elbow detection method (Algorithm S2) and (2) the number of PCs chosen via the BE algorithm (Algorithm S3; the default parameters are used). The BE algorithm is a permutation-based approach for choosing *K* in PCA. Intuitively, it retains PCs that explain more variance in the data than by random chance and discards those that do not. Hence, based on the statistical interpretation of the BE algorithm and the scree plots (examples shown in Figure 7), we believe that the number of PCs chosen via BE should be considered an upper bound of the reasonable number of PCs to choose in GTEx eQTL data [10].

**Figure 7:**
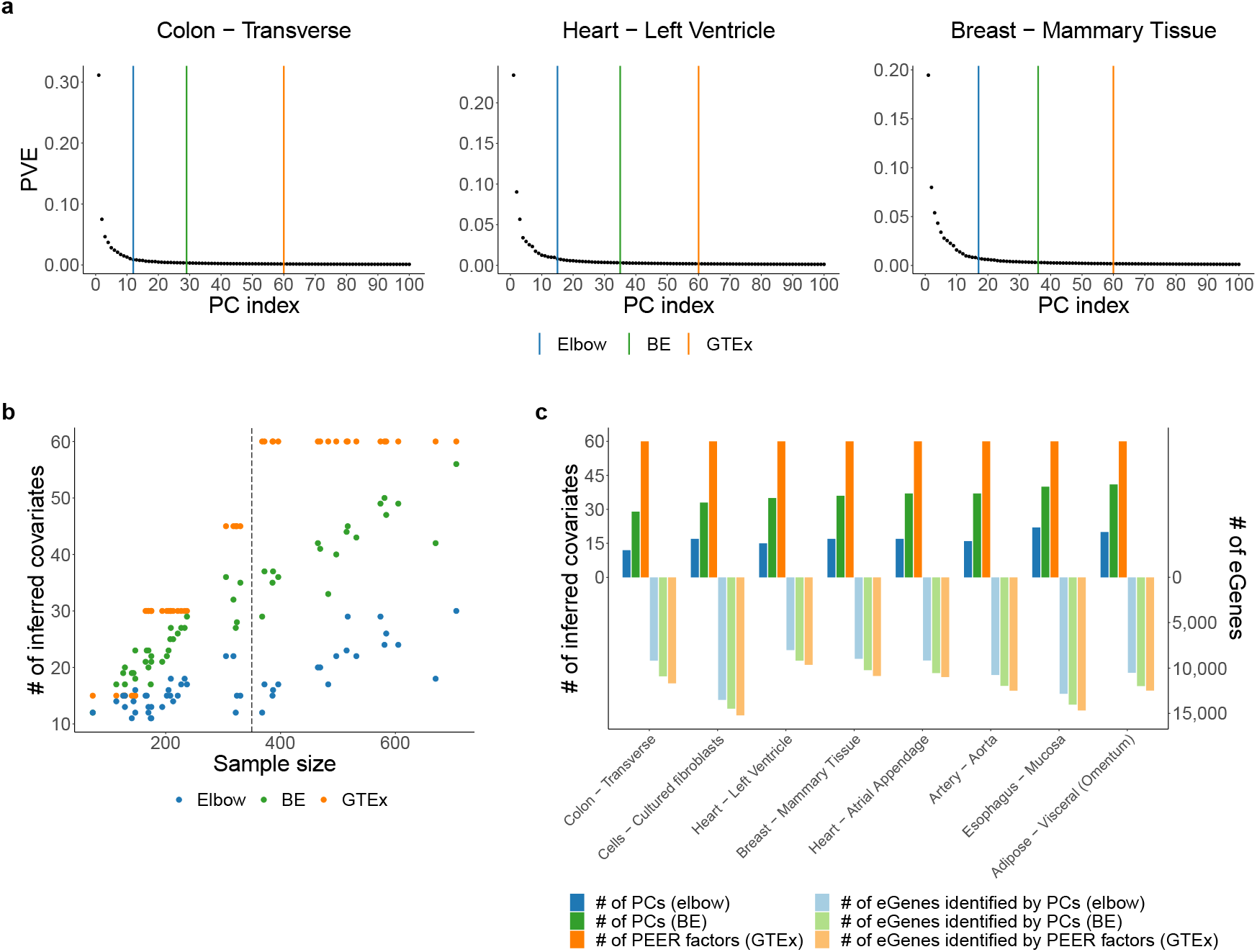
PCA provides insight into the choice of *K*. Recall from Section 2.3 that PEER factors are almost identical to PCs in GTEx eQTL data [10]. Therefore, for each tissue type, we compare the number of PEER factors selected by GTEx to (1) the number of PCs chosen via an automatic elbow detection method (Algorithm S2) and (2) the number of PCs chosen via the BE algorithm (Algorithm S3; the default parameters are used). **a** Example scree plots. **b** This scatter plot contains 49 dots of each color, corresponding to the 49 tissue types with GTEx eQTL analyses. The number of PEER factors selected by GTEx far exceeds the number of PCs chosen via *BE* for many tissue types with sample size above 350 (dashed line), suggesting that the number of PEER factors selected by GTEx may be too large. **c** For the eight tissue types with the largest absolute differences between the number of PEER factors chosen by GTEx and the number of PCs chosen via *BE* (all eight tissue types have sample size above 350), we replace the PEER factors with smaller numbers of PCs in GTEx’s FastQTL pipeline [10, 13] and find that we can reduce the number of inferred covariates to between 20% (12*/*60 = 20%, Colon - Transverse) and 40% (22*/*60 ≈ 36.67%, Esophagus - Mucosa) of the number of PEER factors selected by GTEx without significantly reducing the number of discovered cis-eGenes.

Our results show that the number of PEER factors selected by GTEx is almost always greater than the number of PCs chosen via *BE*, which in turn is almost always greater than the number of PCs chosen via *elbow* (Figure 7). In particular, the number of PEER factors selected by GTEx far exceeds the number of PCs chosen via *BE* for many tissue types with sample size above 350, suggesting that the number of PEER factors selected by GTEx may be too large. This hypothesis is further supported by the fact that we can reduce the number of inferred covariates to between 20% and 40% of the number of PEER factors selected by GTEx without significantly reducing the number of discovered cis-eGenes (Figure 7).

## 3 Discussion

Hidden variable inference is widely practiced as an important step in QTL mapping for improving the power of QTL identification. Popular hidden variable inference methods include SVA, PEER, and HCP. In this work, we show that PCA not only underlies the statistical methodology behind the popular methods (Section 2.4) but is also orders of magnitude faster, better-performing, and much easier to interpret and use (Figure 1; relatedly, Malik and Michoel [48] have pointed out issues with the optimization algorithm used in PANAMA [49]—a variant of PEER, and the computational efficiency of PCA has been reported in other settings, including genomic selection [50]). Our conclusions are consistent with those from Cuomo et al. [51], who conclude that PCA is superior to alternative hidden variable inference methods for improving the power of single-cell eQTL studies.

On the simulation front, we compare the runtime and performance of PCA, SVA, PEER, and HCP via two simulation studies (Section 2.1). In the first simulation study, we follow the data simulation in Stegle et al. [24], the original PEER publication, while addressing its data analysis and overall design limitations. In the second simulation study, we further address the data simulation limitations of Stegle et al. [24] by simulating the data in a more realistic and comprehensive way. Both simulation studies unanimously show that PCA is faster and better-performing. Further, they show that running PEER *with* the known covariates has no advantage over running PEER *without* the known covariates—in fact, running PEER *with* the known covariates makes PEER significantly slower (Figure S6)—and that contrary to claims in Stegle et al. [24, 32], the performance of PEER *does* deteriorate as the number of PEER factors increases (Section 2.1). One caveat of our simulation studies, though, is that the genotype and covariates all have linear effects on the gene expression levels (consistent with Stegle et al. [24] and Wang et al. [41]). But since PCA, SVA, PEER, and HCP are all linear methods or assume linearity (Section 2.4), and so does linear regression, we do not believe our conclusions would change qualitatively if we simulated the data in a nonlinear fashion.

On the real data front, we examine the most recent GTEx eQTL and sQTL data [10] and the 3^′^aQTL data prepared by Li et al. [11] from GTEx RNA-seq reads [9]. While the exact data analysis pipelines are different (Table S3), these studies all choose PEER as their hidden variable inference method (due to lack of data availability, we do not examine more real data sets). Our analysis shows that PEER, the most popular hidden variable inference method for QTL mapping currently, produces identical results as PCA at best (Section 2.3), is at least three orders of magnitude slower than PCA (Figure S6), and can be full of pitfalls. Specifically, we show that in certain cases, PEER factors can be highly correlated with each other and thus fail to capture important variance components of the molecular phenotype data, leading to potential loss of power in QTL identification (Section 2.2). Further, we show from the perspective of PCA that choosing the number of PEER factors by maximizing the number of discoveries (a common approach used by practitioners) may yield inappropriate choices of *K*, leading to model overfit and potential loss of power and precision (Section 2.5).

Between the two PCA approaches, PCA direct (running PCA on the fully processed molecular phenotype matrix *directly* and filtering out the known covariates that are captured well by the top PCs afterwards) and PCA resid (running PCA after regressing out the effects of the known covariates from the molecular phenotype matrix) (Table 1; Section S4), we recommend PCA direct because the two approaches perform similarly in our simulation studies and PCA direct is simpler. In addition, PCA direct can better hedge against the possibility that the known covariates are not actually important confounders because in PCA direct, the known covariates do not affect the calculation of the PCs. We also advise the users to make sure to center and scale their data when running PCA unless they are experts and have a good reason not to.

In addition to the benefits discussed so far, using PCA rather than SVA, PEER, or HCP has another conceptual benefit. While SVA, PEER, and HCP are hidden variable inference (i.e., factor discovery) methods, PCA can be interpreted and used as both a *dimension reduction* and a *factor discovery* method. Therefore, PCs of the molecular phenotype data need not be considered inferred covariates; instead, they can be considered a dimension-reduced version of the molecular phenotype data—by including them as covariates, we are controlling for the effect of the overall gene expression profile on the expression level of any individual gene (taking expression phenotypes as an example). With this perspective, including *phenotype* PCs as covariates is analogous to including *genotype* PCs as covariates (which is commonly done to correct for population stratification [9, 10]). This perspective solves the conundrum that inferred covariates such as PEER factors are often difficult to interpret using known technical and biological variables [52].

To help researchers use PCA in their QTL analysis, we provide an R package `PCAForQTL`, which implements highly interpretable methods for choosing the number of PCs (Algorithms S2 and S3), a graphing function, and more, along with a detailed tutorial. Both resources are freely available at https://github.com/heatherjzhou/PCAForQTL [53]. We believe that using PCA rather than SVA, PEER, or HCP will substantially improve and simplify hidden variable inference in QTL mapping as well as increase the transparency and reproducibility of QTL research.

## 4 Methods

### 4.1 Evaluation metrics

Given a simulated data set, we evaluate each of the 15 methods in Table 1 mainly in three ways (when applicable): runtime, AUPRC, and adjusted *R*^2^ measures (including adjusted *R*^2^, reverse adjusted *R*^2^, and concordance score).

First, we record the runtime of the hidden variable inference step (Section S4; not applicable for Ideal and Unadjusted).

Second, we calculate the area under the precision-recall curve (AUPRC) of the QTL result. We use AUPRC rather than the area under the receiver operating characteristic curve (AUROC) because AUPRC is more appropriate for data sets with imbalanced classes (there are far more negatives than positives in our simulated data sets and in QTL settings in general). Since AUPRC measures the trade-off between the true positive rate (i.e., power) and the false discovery rate (i.e., one minus precision), it is a more comprehensive metric than power. However, to contrast the results in Stegle et al. [24], we also compare the powers of the different methods in Simulation Design 1.

Third, for each simulated data set, each method except Ideal and Unadjusted gets an adjusted *R*^2^ score (short as “adjusted *R*^2^”), a reverse adjusted *R*^2^ score (short as “reverse adjusted *R*^2^”), and a concordance score. The adjusted *R*^2^ score summarizes how well the true hidden covariates can be captured by the inferred covariates; the reverse adjusted *R*^2^ score summarizes how well the inferred covariates can be captured by the true hidden covariates (a low score indicates that the inferred covariates are invalid or “meaningless”); lastly, the concordance score is the average of the previous two scores and thus measures the concordance between the true hidden covariates and the inferred covariates. Specifically, given *m* true hidden covariates and *n* inferred covariates, first, we calculate *m* adjusted *R*^2^’s (regressing each true hidden covariate against the inferred covariates) and *n* reverse adjusted *R*^2^’s (regressing each inferred covariate against the true hidden covariates); then, we average the *m* adjusted *R*^2^’s to obtain the adjusted *R*^2^ score and average the *n* reverse adjusted *R*^2^’s to obtain the reverse adjusted *R*^2^ score; finally, we define the concordance score as the average of the adjusted *R*^2^ score and the reverse adjusted *R*^2^ score.

### 4.2 Selection of representative methods for detailed comparison

Here we describe how we select a few representative methods from the 15 methods for detailed comparison in Simulation Design 2 (Table 1). From Figures 2(d) and S3, we see that the two PCA methods perform almost identically, so for simplicity, we select PCA direct screeK. The two SVA methods perform almost identically as well, so we select SVA BE. For PEER, whether the known covariates are inputted when PEER is run has little effect on the AUPRC. Further, we observe that when we use the true *K*, the factor approach outperforms the residual approach, but when we use a large *K*, the residual approach outperforms the factor approach. Therefore, we select PEER withCov trueK factors and PEER withCov largeK residuals as the representative PEER methods. In addition, Ideal, Unadjusted, and HCP trueK are selected.

### 4.3 A numerical example

Here we provide a simple numerical example of QTL analysis with hidden variable inference by summarizing the setup of GTEx’s cis-eQTL analysis for Colon - Transverse [10].

Let *Y* denote the *n* × *p* fully processed gene expression matrix with *n* = 368 samples and *p* = 25, 379 genes. Let *X*_1_ denotes the *n* × *K*_1_ known covariate matrix with *K*_1_ = 8 known covariates, which include the top five genotype PCs, WGS sequencing platform (HiSeq 2000 or HiSeq X), WGS library construction protocol (PCR-based or PCR-free), and donor sex. Let *X*_inferred_ denote the *n* × *K* inferred covariate matrix with *K* = 60 PEER factors, which are obtained by running PEER on *Y* (Table S3). For gene *j, j* = 1, …, *p*, the relevant genotype data is stored in *S* _*j*_, the *n* × *q* _*j*_ genotype matrix, where each column of *S* _*j*_ corresponds to a local common SNP for gene *j*, and *q* _*j*_ is typically under 15, 000.

Given these input data, the nominal pass (the first step) of FastQTL [13], or equivalently, Matrix eQTL [12], performs a linear regression for each gene and each of its local common SNPs. Specifically, for *j* = 1, …, *p, l* = 1 …, *q* _*j*_, the linear regression represented by the following R `lm()` formula is run:

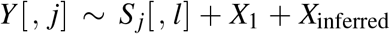

(where *Y* [, *j*] denotes the *j*th column of *Y*, and *S* _*j*_[, *l*] denotes the *l*th column of *S* _*j*_), and the *p*-value for the null hypothesis that the coefficient corresponding to *S* _*j*_[, *l*] is zero (given the covariates) is retained. The top five genotype PCs in *X*_1_ are included in the analysis to correct for population stratification [9, 10] and are typically considered known covariates (see Section 3).

## Availability of data and materials

The R package `PCAForQTL` and a tutorial on using PCA for hidden variable inference in QTL mapping are available at https://github.com/heatherjzhou/PCAForQTL [53]. The code used to generate the results in this work is available at https://doi.org/10.5281/zenodo.6788888 [54]. In addition, this work makes use of the following data and software:

- GTEx V8 public data [10], including fully processed gene expression matrices, fully processed alternative splicing phenotype matrices, known covariates, PEER factors, and QTL results, are downloaded from https://gtexportal.org/home/datasets.
- GTEx V8 protected data [10], specifically, the whole genome sequencing (WGS) phased genotype data, are downloaded from the AnVIL repository with an approved dbGaP application (see https://gtexportal.org/home/protectedDataAccess).
- 3^′^aQTL data prepared by Li et al. [11] from GTEx RNA-seq reads [9] are available from the authors by request.
- SVA R package Version 3.40.0 (https://bioconductor.org/packages/sva/, accessed on October 15th, 2021).
- PEER R package Version 1.3
- (https://bioconda.github.io/recipes/r-peer/README.html, accessed before October 15th, 2021).
- HCP R package Version 1.6 (https://rdrr.io/github/mvaniterson/Rhcpp/, accessed on October 15th, 2021).
- FastQTL (https://github.com/francois-a/fastqtl).

## Authors’ contributions

H.J.Z and J.J.L. conceived the project. H.J.Z. performed the experiments and data analyses and wrote the manuscript. L.L. provided the 3’aQTL data. L.L., Y.L., and W.L. served as advisors. J.J.L. supervised the project. All authors participated in discussions and approved the final manuscript.

## Acknowledgments

The authors would like to thank former and current members of Junction of Statistics and Biology at UCLA for their valuable insight and suggestions, including Elvis Han Cui, Kexin Li, Dr. Xinzhou Ge, Dr. Ruochen Jiang, and Dr. Yiling Chen. The authors would also like to thank the reviewers of this manuscript, including Dr. Tom Michoel, for their insightful comments and suggestions.

## Declarations

### Ethics approval and consent to participate

Not applicable.

### Consent for publication

Not applicable.

## Supplementary materials

### S1 Limitations of the original PEER simulation study

The simulation study in the original PEER publication [24] is limited. We categorize its limitations into three categories: (1) data analysis limitations, (2) overall design limitations, and (3) data simulation limitations.

The data analysis limitations include:

a. The study only compares PEER against the other methods in terms of power, not in terms of false positive rate or false discovery rate (see Section 4.1 for our evaluation metrics).
b. The study does not use PCA or SVA properly (we do; Section S4).
c. The study does not evaluate the different ways of using PEER (we do; Section S4).
d. The study uses ad hoc priors for PEER that are different from the default priors (we use the default priors; Section S4).

The overall design limitations include:

a. The study only simulates one replicate of one experiment. That is, the entire simulation study is based on one simulated data set (we simulate 10 replicates in our first simulation study; Section S2).

The data simulation limitations include (see Table S1 for our solutions):

a. The data dimensions are minimal, with *q* = 100 SNPs in the entire genome.
b. The SNP genotypes are simulated independently and identically with a target minor allele frequency (MAF) of 0.4, so there is no linkage disequilibrium (LD) and a higher average MAF than in real data (the average MAF in GTEx data [10], after SNPs with MAF under 0.01 are filtered out, is about 0.15; Section S3.1).
c. The gene expression levels are primarily driven by trans-regulatory effects rather than cis-regulatory effects or covariate effects (Table S2), inconsistent with the common belief that trans-regulatory effects are generally weaker than cis-regulatory effects.

In addition, the simulation study in the original PEER publication [24] is imperfect in that the description of the data simulation and analysis is vague and inconsistent, and there is no reproducible code. In contrast, we describe our data simulation and analysis in detail (Sections S2 to S4) and provide the code we use to generate the results (see Availability of data and materials).

### S2 Simulation Design 1

#### S2.1 Data simulation

In Simulation Design 1, we perform 10 replicates of the same experiment, where in each replicate, we follow the data simulation in Stegle et al. [24] as closely as possible.

In each replicate, we simulate a data set with *n* = 80 individuals, *p* = 400 genes, *q* = 100 SNPs in the entire genome, *K*_1_ = 3 known covariates, and *K*_2_ = 7 hidden covariates. Let *i, j, l*, and *k* be the indices of individuals, genes, SNPs, and covariates, respectively. That is, *i* = 1, …, *n*; *j* = 1, …, *p*; *l* = 1, …, *q*; and *k* = 1, …, (*K*_1_ + *K*_2_). The data simulation consists of three steps.

In the first step, we simulate *Y*_beforeDSE_, the gene expression matrix before downstream effect, based on

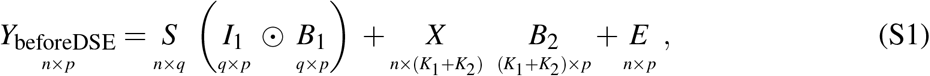

where ⊙ denotes element-wise multiplication. Specifically, in the genotype component, we have

- *S*: genotype matrix. Each entry is drawn independently from *Binom* (2, prob = 0.4). That is, the target MAF is 0.4. In this work, all random sampling is independent unless otherwise specified.
- *I*_1_: effect indicator matrix. Each entry is drawn from *Ber* (0.01).
- *B*_1_: effect size matrix. Each entry is drawn from *N* (0, var = 4).

In the covariate component, we have

- *X* : covariate matrix. Each entry is drawn from *N* (0, var = 0.6). The first *K*_1_ columns are designated as the known covariates (*X*_1_, *n* × *K*_1_), and the last *K*_2_ columns are designated as the hidden covariates (*X*_2_, *n* × *K*_2_).
- *B*_2_: effect size matrix. First, we draw 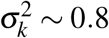 (Γ(shape = 2.5, rate = 0.6))^2^, the covariate-specific effect size variance. Then, we draw 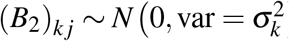.

Lastly, in the noise component, we have

- *E*: noise matrix. First, we draw *τ*_*j*_ ∼ Γ(shape = 3, rate = 1), the gene-specific noise precision. Then, we draw (*E*)_*i j*_ ∼ *N* 0, var = 1*/τ*_*j*_.

In the second step, we simulate *Y*_DSE_, the gene expression matrix due to the downstream effect of genes, based on

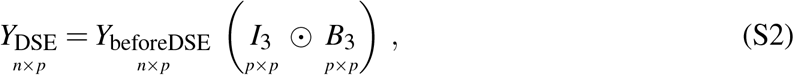

where we have

- *I*_3_: effect indicator matrix. To simulate *I*_3_, we start with a zero matrix. Then, we randomly choose three rows corresponding to genes with at least one cis-QTL (Section S2.2). For each of these three rows, we randomly assign 30 entries corresponding to genes other than the current gene in consideration (avoiding self-loops) to be one.
- *B*_3_: effect size matrix. Each entry is drawn from *N* (8, var = 0.8) for “strong downstream effects” [24].

As we see in Section S2.2, the downstream effect of genes induces trans-QTL relations.

In the third and last step, we define *Y*, the final, observed gene expression matrix, as

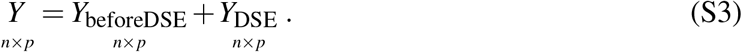

#### S2.2 Definition of truth

In a simulated data set, the cis-QTL relations are encoded in the *q* × *p* binary matrix *I*_1_. The *l j*-th entry being one means that SNP *l* and gene *j* form a cis-QTL pair (i.e., SNP *l* is a cis-QTL for gene *j*).

The trans-QTL relations are encoded in *J*, also a *q* × *p* binary matrix. *J* is defined based on *I*_1_ and *I*_3_. Specifically, SNP *l* and gene *j* form a trans-QTL pair if and only if SNP *l* is a cis-QTL for gene *j*^′^ *and* gene *j*^′^ has downstream effect on gene *j, j*^′^ ≠ *j*.

The overall truth is encoded in 𝟙((*I*_1_+*J*) ≥ 1), again a *q* × *p* binary matrix. We use this matrix as the truth when calculating AUPRCs. The *l j*-th entry being one means that SNP *l* and gene *j* form a cis-QTL or trans-QTL pair (or both).

**Table S1:** Summary of the main differences between Simulation Design 1 and Simulation Design 2. Highlighted in blue are the major data simulation limitations (Section S1) of Simulation Design 1, all of which we address in Simulation Design 2.

|  | Simulation Design 1 | Simulation Design 2 |
| --- | --- | --- |
| Data simulation | Follows Stegle et al. [24] | Loosely based on Wang et al. [41] |
| # of experiments | 1 | 176 |
| # of replicates per experiment | 10 | 2 |
| # of simulated data sets | 10 | 352 |
| Genotype data | Simulated (no LD, high MAF) | Real genotype data from GTEx [10] |
| Cis-QTL relations present | ✓ | ✓ |
| Trans-QTL relations present | ✓ | ✗ |
| Source(s) of expression variation | Primarily trans-regulatory effects | Carefully controlled genotype effects and covariate effects |
| # of individuals | $n = 80$ | $n = 838$ |
| # of genes | $p = 400$ | $p = 1,000$ |
| # of SNPs | $q = 100$ SNPs in the entire genome | $q = 1,000$ local common SNPs per gene |
| # of known covariates | $K_1 = 3$ | $K_1 = 2, 3, 5, \text{ or } 8$ depending on the experiment |
| # of hidden covariates | $K_2 = 7$ | $K_2 = 3, 7, 15, \text{ or } 22$ depending on the experiment |

**Table S2:** In Simulation Design 1, which follows the data simulation in Stegle et al. [24] as closely as possible, the gene expression levels are primarily driven by trans-regulatory effects rather than cis-regulatory effects or covariate effects. Var (*Y*_beforeDSE_) is defined as the variance of the *n* × *p* entries of *Y*_beforeDSE_, and the other variances in the table are defined similarly. We find that Var (*Y*_DSE_) */* Var (*Y*) is above 90% on average.

| Replicate | $\text{Var}(Y_{\text{beforeDSE}})$ | $\text{Var}(Y_{\text{DSE}})$ | $\text{Var}(Y)$ | $\text{Var}(Y_{\text{beforeDSE}}) / \text{Var}(Y)$ | $\text{Var}(Y_{\text{DSE}}) / \text{Var}(Y)$ |
| --- | --- | --- | --- | --- | --- |
| 1 | 124.34 | 1757.07 | 1889.57 | 6.58% | 92.99% |
| 2 | 140.92 | 1505.17 | 1677.64 | 8.40% | 89.72% |
| 3 | 213.56 | 929.96 | 1169.18 | 18.27% | 79.54% |
| 4 | 71.85 | 761.01 | 855.39 | 8.40% | 88.97% |
| 5 | 123.07 | 2434.45 | 2574.51 | 4.78% | 94.56% |
| 6 | 74.94 | 1029.29 | 1092.65 | 6.86% | 94.20% |
| 7 | 148.61 | 2490.72 | 2628.93 | 5.65% | 94.74% |
| 8 | 79.36 | 796.55 | 868.05 | 9.14% | 91.76% |
| 9 | 54.62 | 1340.10 | 1390.72 | 3.93% | 96.36% |
| 10 | 65.64 | 831.89 | 895.90 | 7.33% | 92.86% |
| Average |  |  |  | 7.93% | **91.57%** |

**Figure S1:**
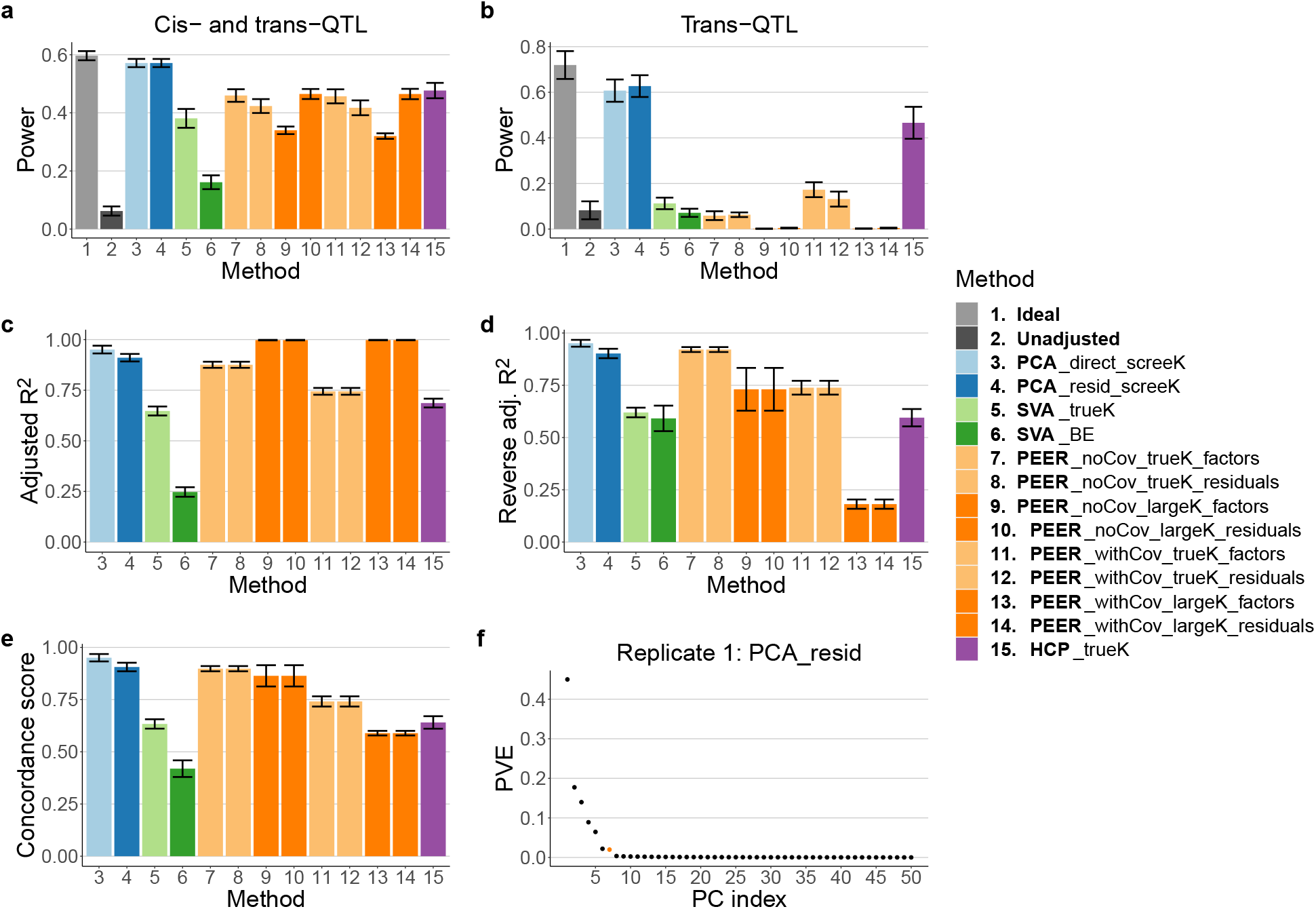
Comparison of all 15 methods (Table 1) in terms of power and adjusted *R*^2^ measures in Simulation Design 1 (the height of each bar represents the average across simulated data sets) and an example scree plot. **a, b** PCA is more powerful than SVA, PEER, and HCP both when we consider all QTL relations (**a**) and when we focus on trans-QTL relations (**b**). Binary decisions are made based on *p*-values using the Benjamini-Hochberg (BH) procedure and a target false discovery rate of 0.05. **c, d, e** PCA performs the best in terms of concordance score. PEER with a large *K* (dark orange bars) performs well in terms of adjusted *R*^2^ but less well in terms of reverse adjusted *R*^2^. **f** An example scree plot that unambiguously suggests the true number of hidden covariates, seven in this case, as the reasonable number of PCs to choose (the *y*-axis represents the proportion of variance explained).

### S3 Simulation Design 2

#### S3.1 Data simulation

In Simulation Design 2, we use real genotype data from GTEx [10], focus on cis-QTL detection, and carefully control the genotype effects and covariate effects in 176 experiments with two replicates per experiment. This simulation design takes inspiration from and is loosely based on Wang et al. [41].

In each experiment-replicate combination, we simulate a data set with *n* = 838 individuals, *p* = 1, 000 genes, *q* = 1, 000 local common SNPs per gene, *K*_1_ known covariates, and *K*_2_ hidden covariates (the values of *K*_1_ and *K*_2_ depend on the experiment; see below). Again, let *i, j, l*, and *k* be the indices of individuals, genes, SNPs, and covariates, respectively. That is, *i* = 1, …, *n*; *j* = 1, …, *p*; *l* = 1, …, *q*; and *k* = 1, …, (*K*_1_ + *K*_2_).

We begin by obtaining *SArray*, the *n* × *q* × *p* genotype array that remains constant throughout Simulation Design 2. *SArray*[,, *j*], an *n* × *q* matrix, is the genotype matrix for the *q* local common SNPs for gene *j*. We obtain *SArray* with the following steps:

a. Download the whole genome sequencing (WGS) phased genotype data for *n* = 838 individuals from GTEx V8 [10].
b. Randomly select *p* = 1, 000 genes from the more than 20,000 genes on chromosomes 1 to 22.
c. For each gene, obtain the genotype data for the *q* = 1, 000 SNPs with MAF≥ 0.01 that are the closest to the gene’s transcription start site (TSS); we find that these SNPs are almost always within 1 Mb of the TSS. The average MAF of *SArray*, calculated as 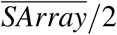, the average of all entries of *SArray* divided by 2, is 0.1474385 ≈ 0.15.

Each experiment is characterized by four attributes:

a. Number of effect SNPs per gene (`numOfEffectSNPs`): 1 or `random`.
b. Number of covariates (`numOfCovariates`): 5, 10, 20, or 30.
  - Number of known covariates (*K*_1_): 2, 3, 5, 8, respectively.
  - Number of hidden covariates (*K*_2_): 3, 7, 15, 22, respectively.
c. Proportion of variance explained by genotype (`PVEGenotype`): 0.05, 0.1, 0.2, or 0.3.
d. Proportion of variance explained by covariates (`PVECovariates`): minimum 0.3, maximum 1 − 0.05 − `PVEGenotype`, in increments of 0.1. For example, when `PVEGenotype` = 0.05, `PVECovariates` takes seven possible values: 0.3, 0.4, 0.5, …, 0.9.

Therefore, we have a total of 2 × 4 ×(7 + 6 + 5 + 4) = 8 × 22 = 176 experiments covering typical scenarios in QTL studies [41]. Following Wang et al. [41], we use the term “effect SNPs” to refer to SNPs that have a nonzero cis effect on a given gene.

Given `numOfEffectSNPs`, `numOfCovariates`, `PVEGenotype`, and `PVECovariates`, we simulate each data set based on

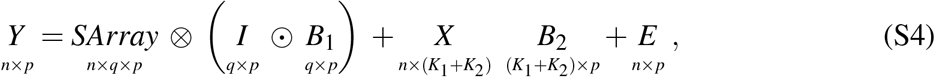

where *Y* is the gene expression matrix, and ⊗ is defined as

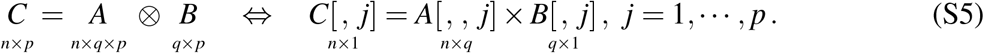

Specifically, in the genotype component, we have

- *SArray*: genotype array. *SArray*[,, *j*], an *n* × *q* matrix, is the genotype matrix for the *q* local common SNPs for gene *j* (see above).
- *I*: effect indicator matrix.
  – If `numOfEffectSNPs` = 1, then for each column, we randomly assign one entry to be one while keeping the other entries zero.
  – If `numOfEffectSNPs` = `random`, then each entry of *I* is drawn from *Ber* (1*/q*). This means that for each gene, the number of effect SNPs is drawn from *Binom* (*q*, prob = 1*/q*). This binomial distribution approximates the empirical distribution of the number of independent cis-eQTLs per gene in GTEx data [10] well (Figure S2).
- *B*_1_: effect size matrix. Each entry is drawn from *N* (0, 1).

In the covariate component, we have

- *X* : covariate matrix. Each entry is drawn from *N* (0, 1). As in Simulation Design 1, the first *K*_1_ columns are designated as the known covariates (*X*_1_, *n* × *K*_1_), and the last *K*_2_ columns are designated as the hidden covariates (*X*_2_, *n* × *K*_2_).
- *B*_2_: effect size matrix. Each entry is drawn from *N* (0, 1) and scaled (see below).

Lastly, in the noise component, we have

- *E*: noise matrix. Each entry is drawn from *N* (0, 1) and scaled (see below).

Alternatively, (S4) can be written as

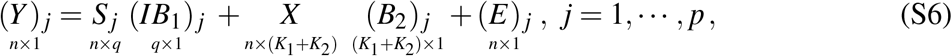

where (*Y*) _*j*_, (*IB*_1_) _*j*_, (*B*_2_) _*j*_, and (*E*) _*j*_ denote the *j*th column of *Y, I* ⊙ *B*_1_, *B*_2_, and *E*, respectively, and *S* _*j*_ denotes *SArray*[,, *j*].

The scaling for *B*_2_ and *E* is to ensure that `PVEGenotype` and `PVECovariates` are as desired. Specifically, for gene *j*, if Var *S* _*j*_ (*IB*_1_) _*j*_ ≠ 0, then we scale (*B*_2_) _*j*_ so that

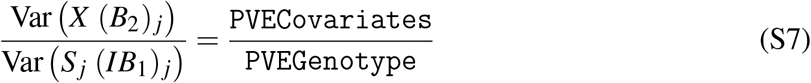

and separately scale (*E*) _*j*_ so that

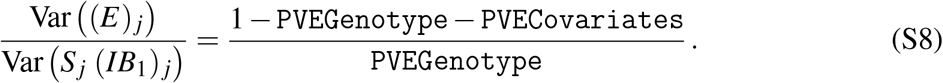

If Var (*S* _*j*_ (*IB*_1_) _*j*_) = 0 (which is the case when (*IB*_1_) _*j*_ is a zero vector, i.e., when gene *j* has zero effect SNPs), then we only scale (*E*) _*j*_ so that

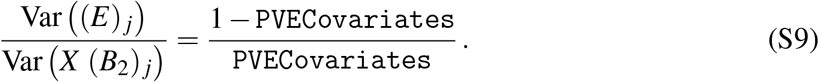

#### S3.2 Definition of truth

In a simulated data set, *I* is a *q* × *p* binary matrix. The *l j*-th entry being one means that the *l*th local common SNP for gene *j* is an effect SNP for gene *j*. However, due to LD, the expression level of a gene may be strongly associated with SNPs other than its effect SNPs.

Therefore, we define *I*_cor_, also a *q* × *p* binary matrix, based on *SArray* and *I* and use it as the truth when calculating AUPRCs. The *l j*-th entry of *I*_cor_ is defined as one if and only if the *l*th local common SNP for gene *j* is highly correlated with *any* of gene *j*’s effect SNPs (correlation ≥ 0.9 in absolute value).

**Figure S2:**
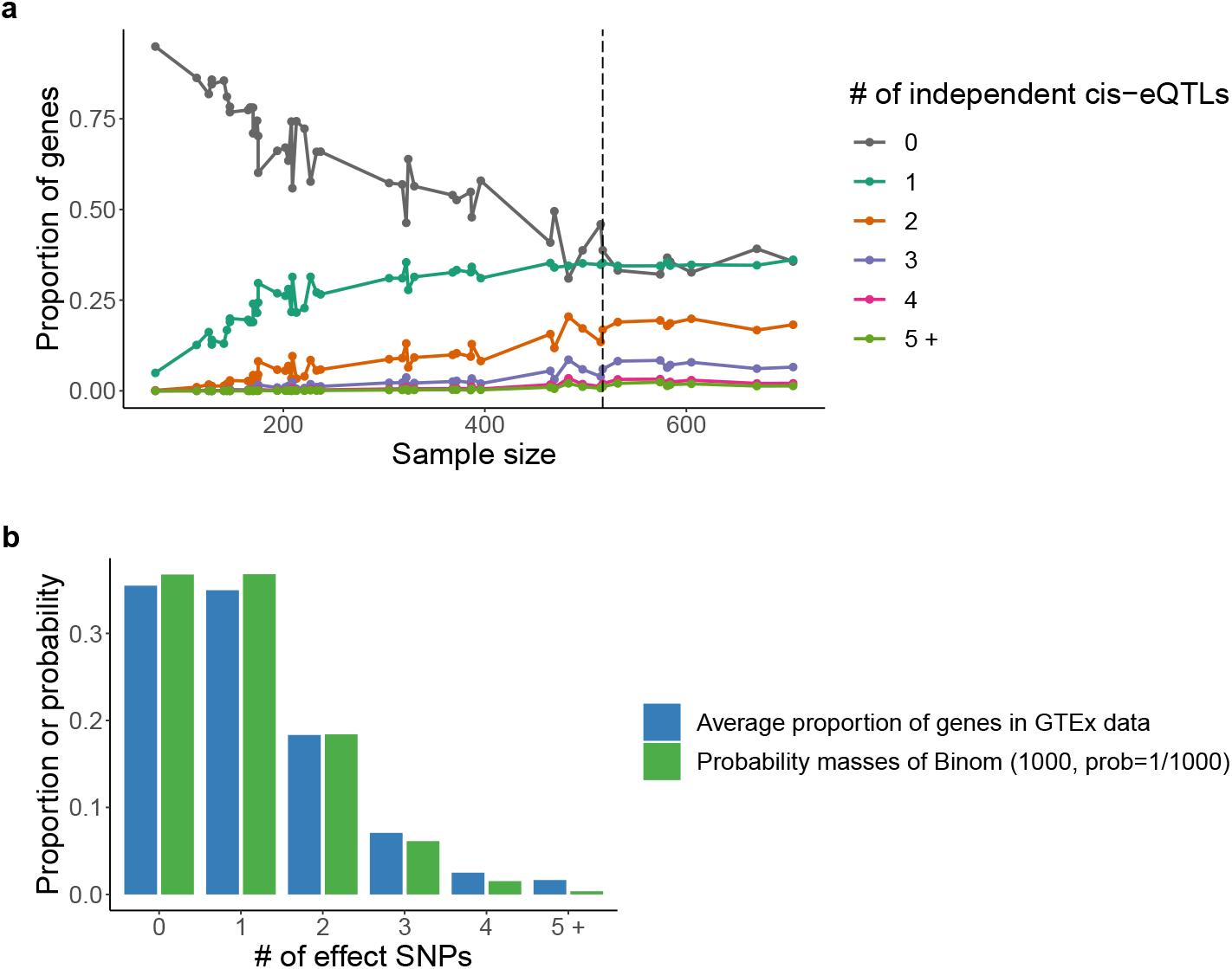
In Simulation Design 2, we find that *Binom* (1000, prob = 1*/*1000) approximates the empirical distribution of the number of independent cis-eQTLs per gene in GTEx data [10] well. **a** Given a tissue type, which corresponds to a sample size, we plot the proportion of genes with 0, 1, 2, 3, 4, or 5 or more independent cis-eQTLs (the proportions add up to one; data from GTEx [10]). We find that the proportions stabilize once the sample size reaches about 517 (dashed line). **b** For the eight tissue types with sample size ≥ 517, we take the average proportion of genes with 0 independent cis-eQTLs, 1 independent cis-eQTL, etc. and plot them in the blue bars. The green bars represent the probability mass function of *Binom* (1000, prob = 1*/*1000) (with the tail probabilities combined together).

**Figure S3:**
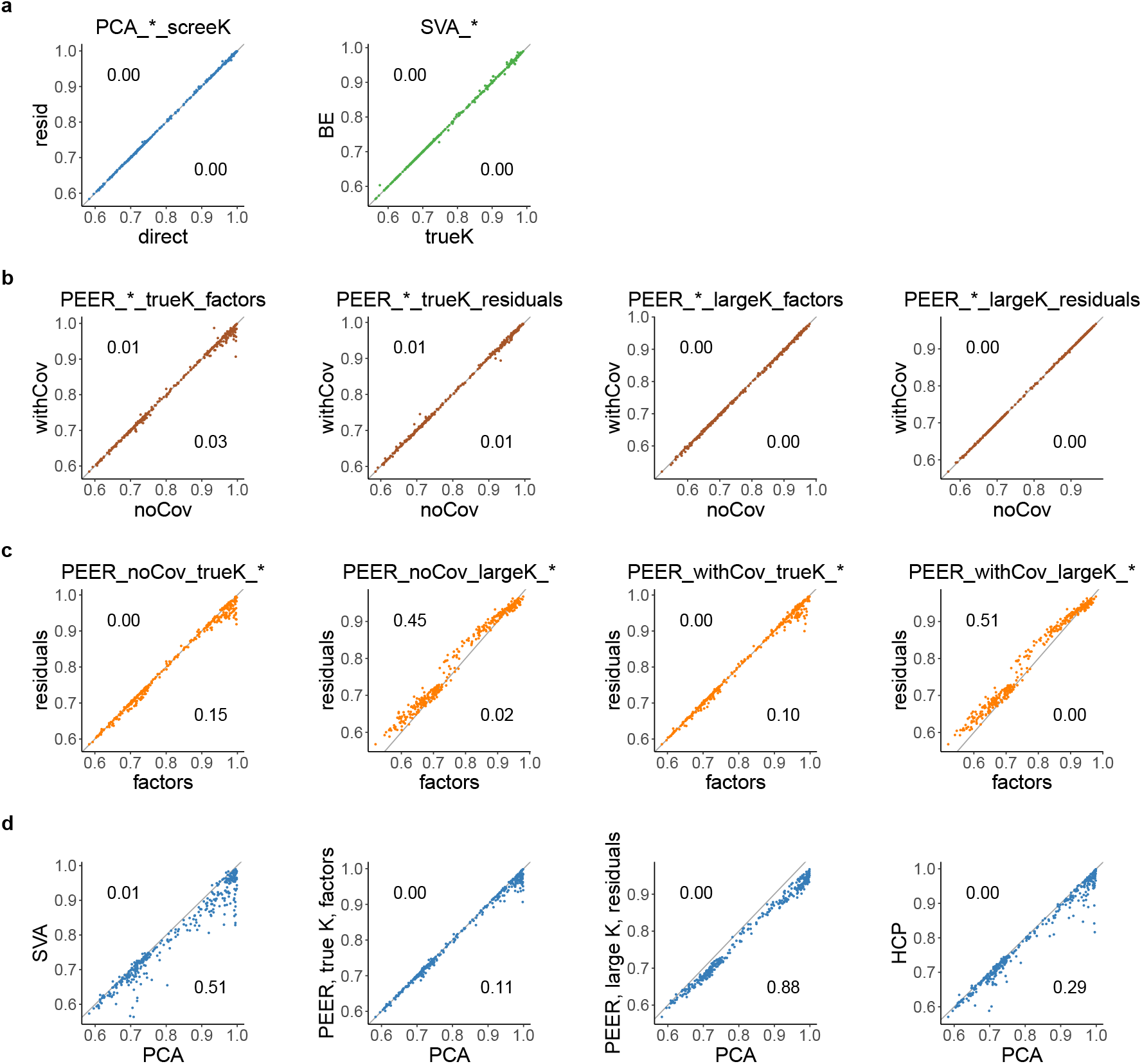
This figure shows how we select a few representative methods from the 15 methods for detailed comparison in Simulation Design 2 (**a, b, c**) and a dataset-by-dataset comparison of the selected representative methods (**d**). The *x*-axis and *y*-axis both represent AUPRCs of different methods. Each scatter plot contains 352 points, each of which corresponds to a simulated data set in Simulation Design 2. The number on the upper-left corner of each scatter plot represents the proportion of points that satisfy *y >* 1.02 *x*, and the number on the lower-right corner represents the proportion of points that satisfy *x >* 1.02 *y*, where *x* and *y* denote the coordinates of each point. **a** The two PCA methods perform almost identically, so for simplicity, we select PCA direct screeK. The two SVA methods perform almost identically as well, so we select SVA BE. **b** Whether the known covariates are inputted when PEER is run has little effect on the AUPRC. **c** When we use the true *K*, the factor approach outperforms the residual approach, but when we use a large *K*, the residual approach outperforms the factor approach. Therefore, we select PEER withCov trueK factors and PEER withCov largeK residuals as the representative PEER methods. **d** Among the selected representative methods, PCA outperforms SVA, PEER, and HCP in terms of AUPRC in 11% to 88% of the simulated data sets and underperforms them in close to 0% of the simulated data sets.

**Figure S4:**
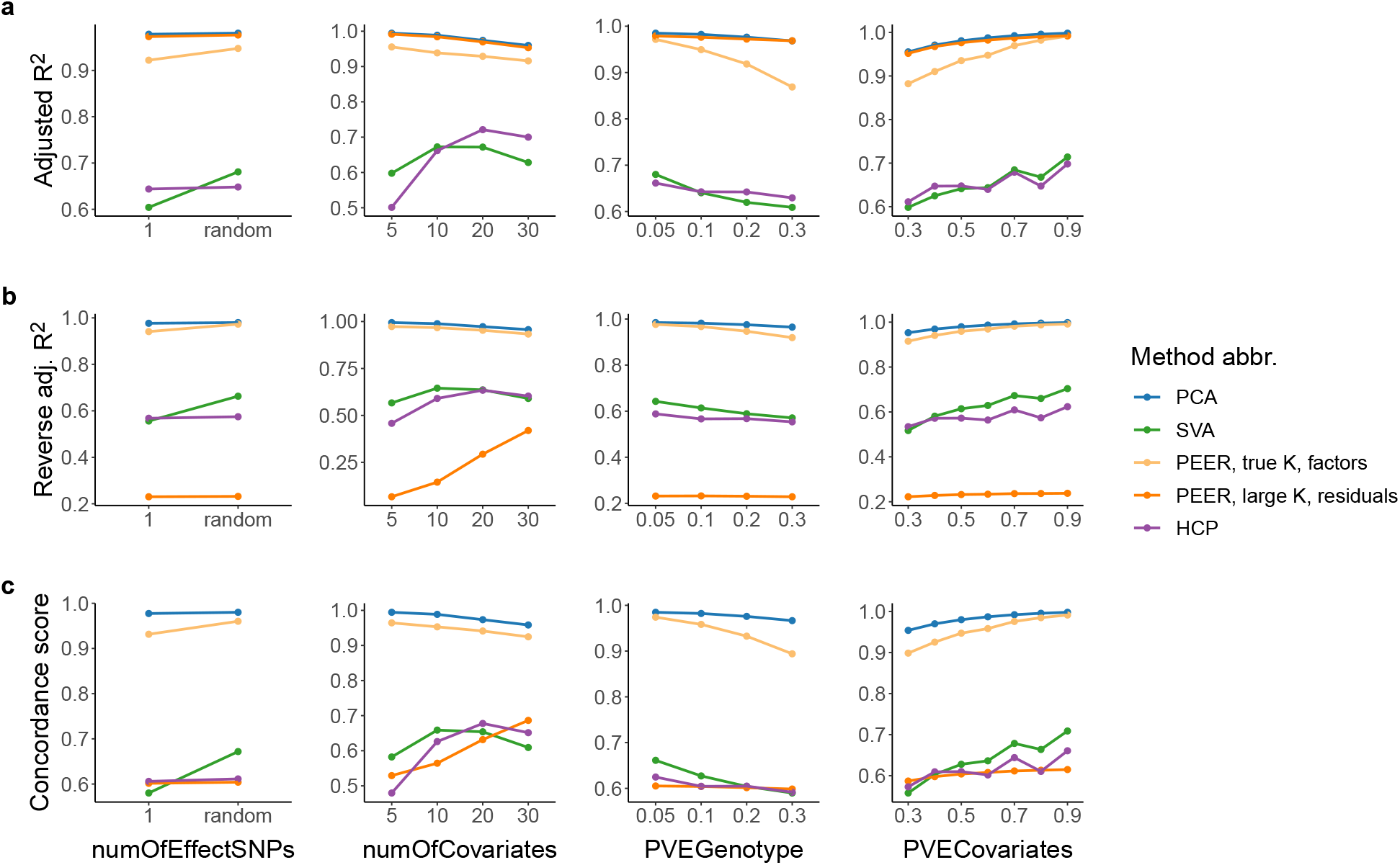
Detailed adjusted *R*^2^, reverse adjusted *R*^2^, and concordance score comparison of the selected representative methods (Table 1) in Simulation Design 2. Each point represents the average across simulated data sets. PCA performs the best in all three regards. PEER with a large *K* (dark orange line) performs well in terms of adjusted *R*^2^ but falls short in terms of reverse adjusted *R*^2^.

### S4 Compared methods

We compare the runtime and performance of 15 methods based on simulation studies, including Ideal, Unadjusted, and 13 variants of PCA, SVA, PEER, and HCP (Table 1). The details of Simulation Design 1 and Simulation Design 2 are described in Sections S2 and S3, respectively. Recall that in each simulated data set, *Y* denotes the gene expression matrix (*n* × *p*, sample by gene), *X*_1_ denotes the known covariate matrix (*n* × *K*_1_, sample by covariate), and *X*_2_ denotes the hidden covariate matrix (*n* × *K*_2_, sample by covariate). The genotype information is stored in *S* in Simulation Design 1 and *SArray* in Simulation Design 2. In this work, we use *K* to denote the number of inferred covariates, which are called PCs, SVs, PEER factors, and HCPs in PCA, SVA, PEER, and HCP, respectively.

Given a simulated data set, each of the 15 methods consists of two steps: hidden variable inference step (not applicable for Ideal and Unadjusted) and QTL step. In the hidden variable inference step, we run PCA, SVA, PEER, or HCP to obtain the inferred covariates (and the expression residuals in the case of PEER; Figure 1). In the QTL step, given a gene-SNP pair, we run a linear regression with the gene expression vector (or the residual vector from PEER) as the *response* and the genotype vector and covariates as *predictors*, where the choice of the response and covariates depends on the method (Table 1); thus we obtain the *p*-value for the null hypothesis that the coefficient corresponding to the genotype vector is zero given the covariates. In Simulation Design 1, we investigate the association between each gene’s expression level and each SNP in the *entire genome* for a simultaneous detection of cis-QTL and trans-QTL relations. In Simulation Design 2, we investigate the association between each gene’s expression level and each of the gene’s *local common SNPs* for a cis-QTL analysis.

For Ideal, we assume that *X*_2_ is known. Therefore, we use *X*_1_ and *X*_2_ as covariates in the QTL step. For Unadjusted, we use *X*_1_ as the covariates.

We devise two ways to use PCA to account for the hidden covariates. For PCA direct screeK, we run PCA on *Y* directly. For PCA resid screeK, we first residualize *Y* against *X*_1_ and then run PCA on the residual matrix. In this work, PCA is run *with* centering and scaling unless otherwise specified; given *A*, an *n* × *p*_1_ matrix, and *B*, an *n* × *p*_2_ matrix, both observation by feature, to *residualize A against B* means to take each column of *A*, regress it against *B*, and replace the original column of *A* with the residuals from the linear regression. For both methods, since the scree plots always suggest the true “number of hidden covariates” (*K*_1_ + *K*_2_ for PCA direct screeK, *K*_2_ for PCA resid screeK) as the reasonable number of PCs to choose within plus or minus one (usually exactly; Figure S1), we set the number of PCs to be the true “number of hidden covariates”. For PCA direct screeK, we filter out the known covariates that are captured well by the top PCs (unadjusted *R*^2^ ≥ 0.9) and use the remaining known covariates along with the top PCs as covariates in the QTL step. For PCA resid screeK, no filtering is needed.

Here we describe the two hidden variable inference methods for SVA: SVA trueK and SVA BE. Since the SVA package [31] requires the user to input at least one variable of interest (Figure 1) and using too many variables of interest causes the package to fail, when running SVA, we input the top PC of the genotype matrix (*S* in Simulation Design 1, collapsed version of *SArray* in Simulation Design 2) as the variable of interest. We also input *X*_1_ as the known covariates because the package documentation indicates that the known covariates should be provided if available. The SVA package allows the user to specify *K*. Alternatively, it can automatically choose *K* using a slightly modified version of the Buja and Eyuboglu (BE) algorithm [44, 47]. Therefore, in SVA trueK, we set *K* = *K*_2_, and in SVA BE, we let the package choose *K* automatically. In both cases, we use *X*_1_ and the surrogate variables (SVs) as covariates in the QTL step.

There are several different ways to use PEER [32] but no consensus in the literature on which one is the best. In the hidden variable inference step, PEER can be run with or without the known covariates when there are known covariates available (Stegle et al. [32] do not give an explicit recommendation as to which approach should be used, and both approaches are used in practice [9–11, 16]), and *K* has to be specified by the user (Stegle et al. [24, 32] claim that the performance of PEER does not deteriorate as *K* increases). In the QTL step, one can include the PEER factors as covariates (we call this the “factor approach”) or use the expression residuals outputted by PEER as the response (and not use any known or inferred covariates; we call this the “residual approach”). For completeness, we compare 2^3^ = 8 ways of using PEER (the default priors are always used): PEER is run with or without the known covariates; PEER is run using the true “number of hidden covariates” (*K*_1_ + *K*_2_ when PEER is run without the known covariates, *K*_2_ when PEER is run with the known covariates) or using a large *K* (*K*=50); and either the factor approach or the residual approach is used in the QTL step.

The HCP package requires the user to specify *K* and three tuning parameters: *λ*_1_, *λ*_2_, and *λ*_3_ (Section S5.2). The package documentation suggests choosing *K* and the tuning parameters via a grid search. However, no specific recommendations are given regarding the choice of the score function. In practice, users of HCP often choose *K* and the tuning parameters by maximizing the number of discoveries [22, 23]. For our simulation studies, such an approach would be computationally prohibitive. Therefore, for simplicity, we set *K* = *K*_2_ and *λ*_1_ = *λ*_2_ = *λ*_3_ = 1; the latter is because we do not want to give more weight to the penalty terms than the main term in the objective function (Section S5.2).

**Table S3:** Summary of QTL analyses performed by GTEx [9, 10] and Li et al. [11]. “INT” in (D) stands for “inverse normal transform” [42]. (E), (F), and (G) summarize how PEER is used (Section S4). GTEx [9, 10] chooses the number of PEER factors for its eQTL analyses (including cis and trans) by maximizing the number of discovered cis-eGenes for each pre-defined sample size bin. The number of PEER factors selected is 15 for *n <* 150, 30 for *n* ∈ [150, 250), and 35 for *n* ≥ 250 for GTEx V6p eQTL analyses [9] and 15 for *n <* 150, 30 for *n* ∈ [150, 250), 45 for *n* ∈ [250, 350), and 60 for *n* ≥ 350 for GTEx V8 eQTL analyses [10], where *n* denotes the sample size. Li et al. [11] use the numbers of PEER factors chosen by GTEx [9].

| Reference | GTEx data version | QTL analysis | Data transformation | Known covariates inputted | # of PEER factors | Factor or residual approach |
| --- | --- | --- | --- | --- | --- | --- |
| (A) | (B) | (C) | (D) | (E) | (F) | (G) |
| GTEx Consortium [9] | V6p | eQTL (cis and trans) | INT within feature | No | Maximizes cis-eGenes | Factor |
| GTEx Consortium [10] | V8 | eQTL (cis and trans) | INT within feature | No | Maximizes cis-eGenes | Factor |
|  |  | sQTL (cis and trans) | INT within sample | No | 15 | Factor |
| Li et al. [11] | V7 | 3'aQTL (cis) | No transformation | Yes | Follows GTEx [9] | Factor |

**Figure S5:**
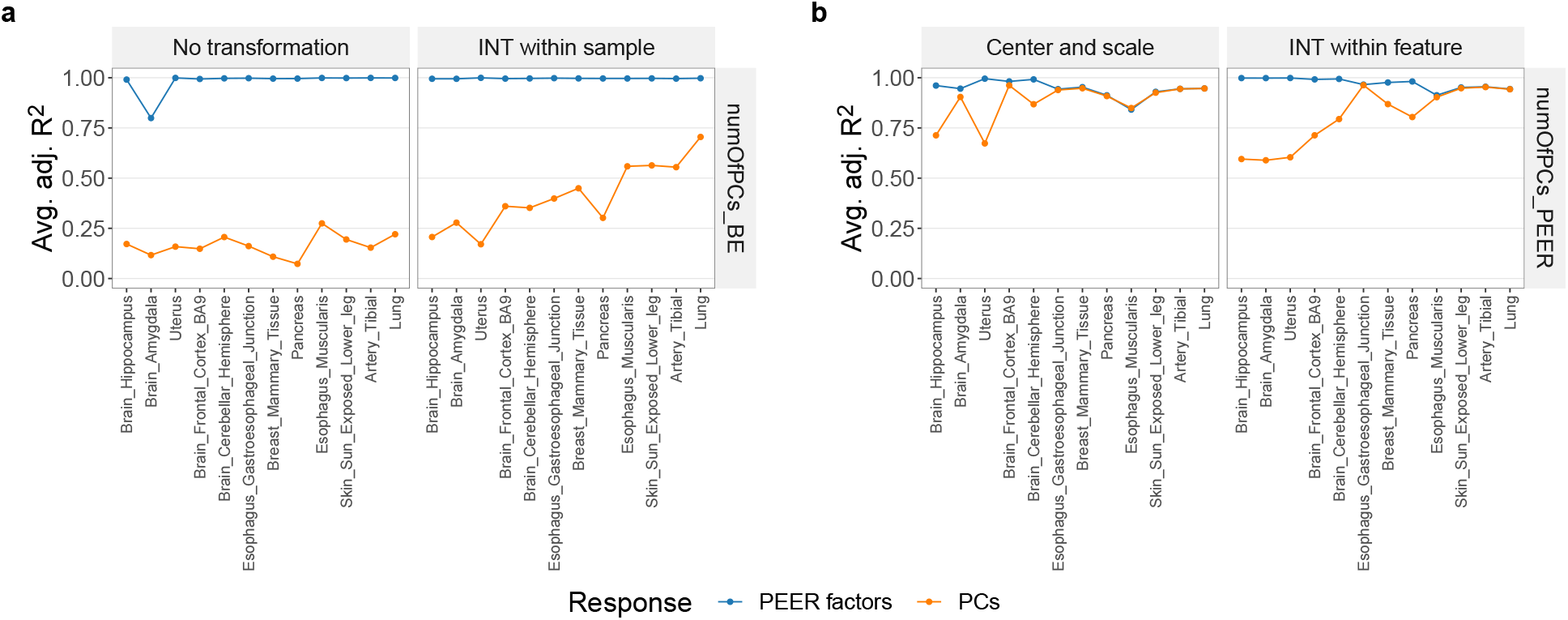
In the 3^′^aQTL data prepared by Li et al. [11] from GTEx RNA-seq reads [9], PEER factors fail to capture important variance components of the molecular phenotype data when the data transformation method is “No transformation” or “INT within sample” (**a**; the numbers of PCs are chosen via BE (Algorithm 3)). On the other hand, PEER factors span roughly the same linear subspace as the top PCs when the data transformation method is “Center and scale” or “INT within feature”, but the top PCs can almost always capture the PEER factors better than the PEER factors can capture the top PCs (**b**; the numbers of PCs are equal to the numbers of PEER factors). Given *m* PEER factors and *n* PCs from the same post-transformation molecular phenotype matrix (*m* ≥ *n* in **a**, *m* = *n* in **b**), we calculate *m* adjusted *R*^2^’s by regressing each PEER factor against the PCs and plot the average in blue. Similarly, we calculate *n* adjusted *R*^2^’s by regressing each PC against the PEER factors and plot the average in orange.

#### Algorithm S1: Reordering of PEER factors based on PCs (Figure 5)

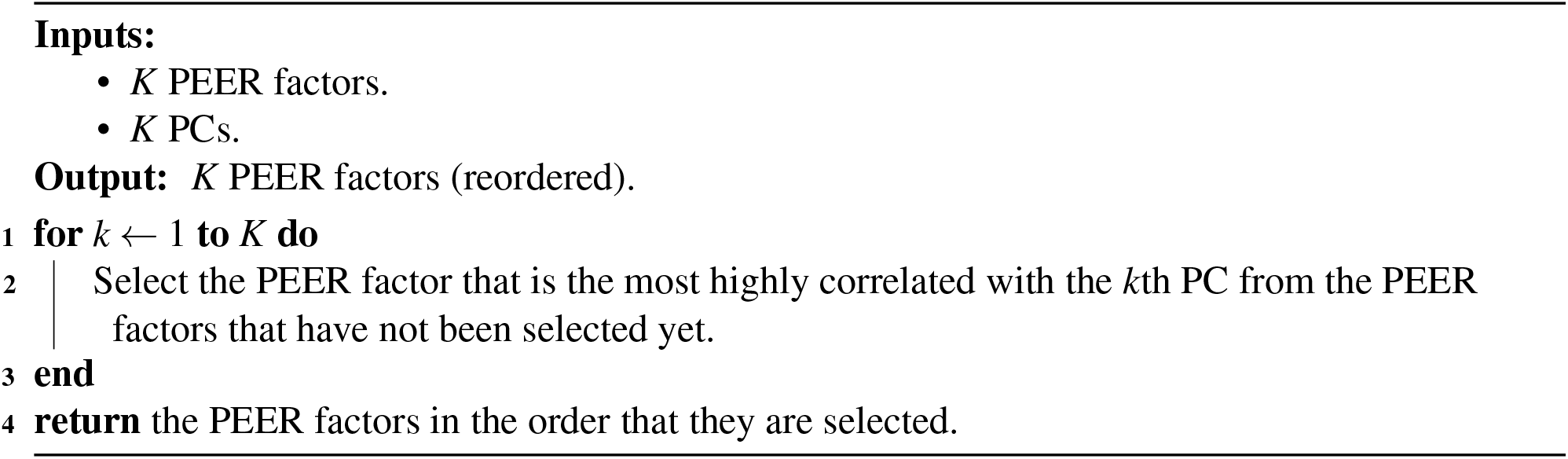

**Figure S6:**
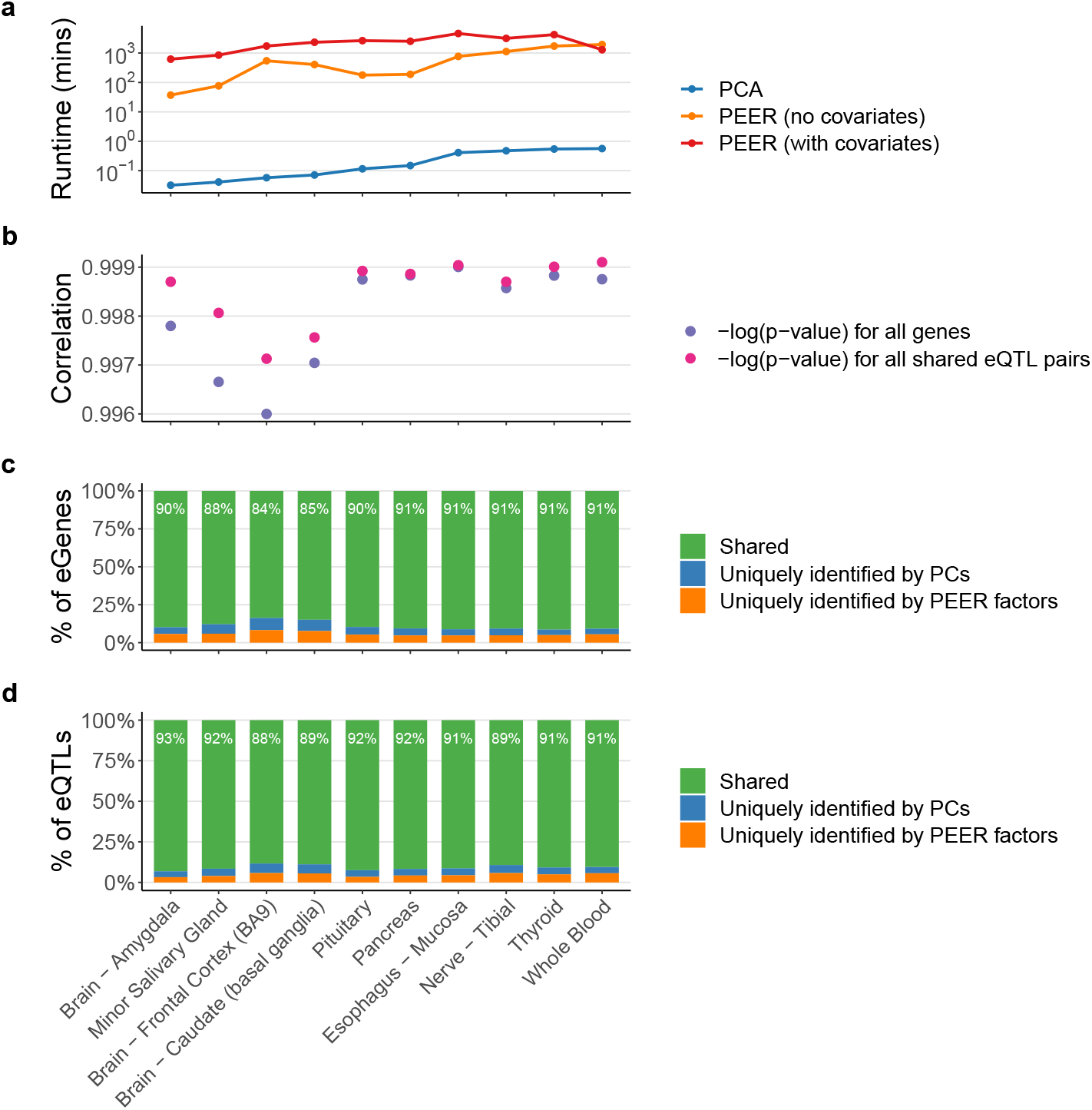
In GTEx eQTL data [10], PEER is at least three orders of magnitude slower than PCA (**a**), and replacing the PEER factors with PCs in GTEx’s FastQTL pipeline [10, 13] does not change the cis-eQTL results much (**b, c, d**). The *x*-axis shows 10 randomly selected tissue types with increasing sample sizes. **a** For a given gene expression matrix, running PEER without the known covariates (GTEx’s approach) takes up to about 1, 900 minutes (equivalent to about 32 hours; Whole Blood), while running PCA (with centering and scaling; our approach) takes no more than a minute. For comparison, we also run PEER *with* the known covariates using the numbers of PEER factors selected by GTEx. This approach takes even longer (up to about 4, 600 minutes, equivalent to about 77 hours; Esophagus - Mucosa). **b** The *p*-values produced by GTEx’s approach and our approach are highly correlated (correlations between the negative common logarithms are shown). **c, d** The overlap of the identified eGenes and eQTL pairs between the two approaches is generally around 90% (see Figure S7 for more detail).

**Figure S7:**
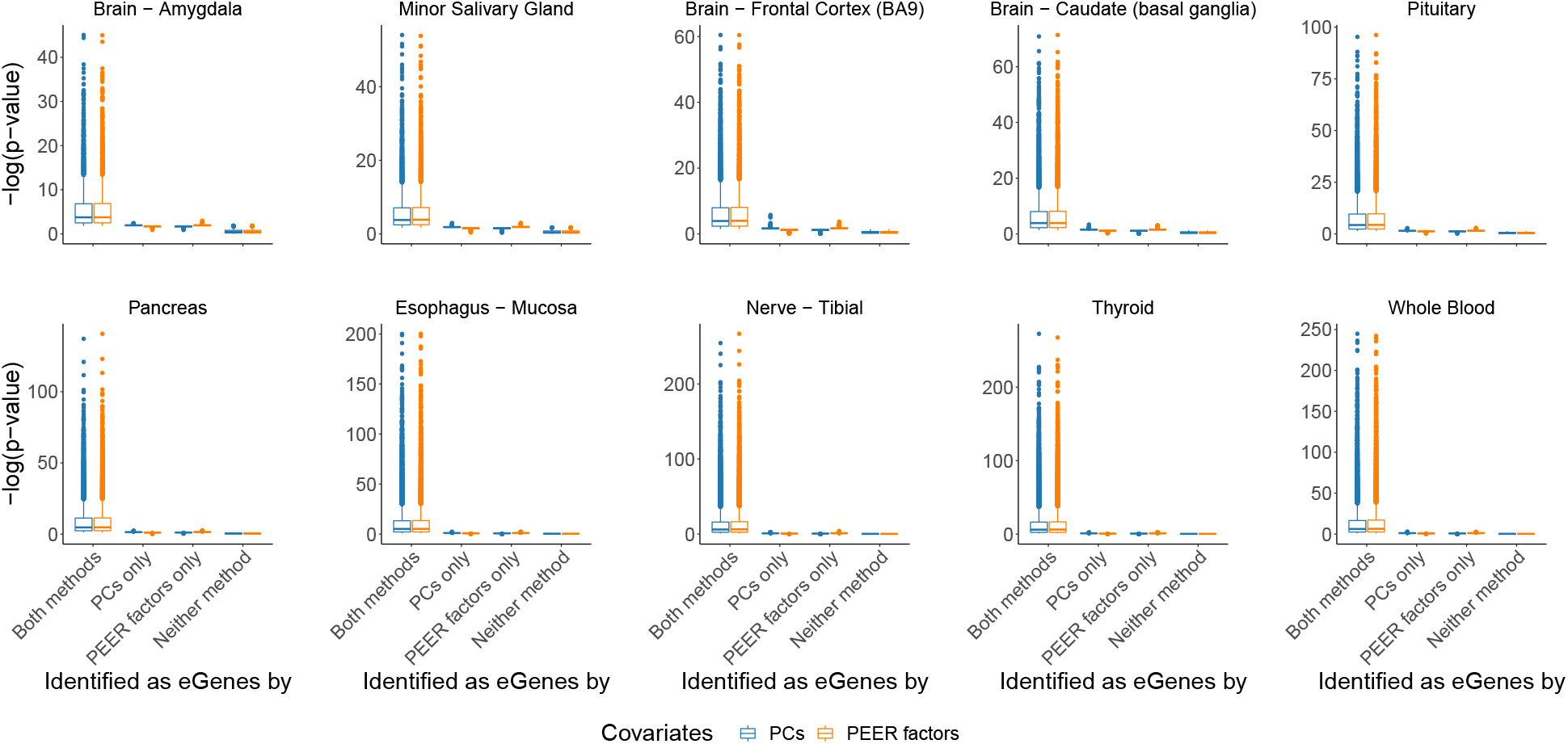
Following the analysis in Figure S6, we find that the eGenes uniquely identified by PCs or PEER factors have marginal *p*-values compared to those identified by both methods.

**Figure S8:**
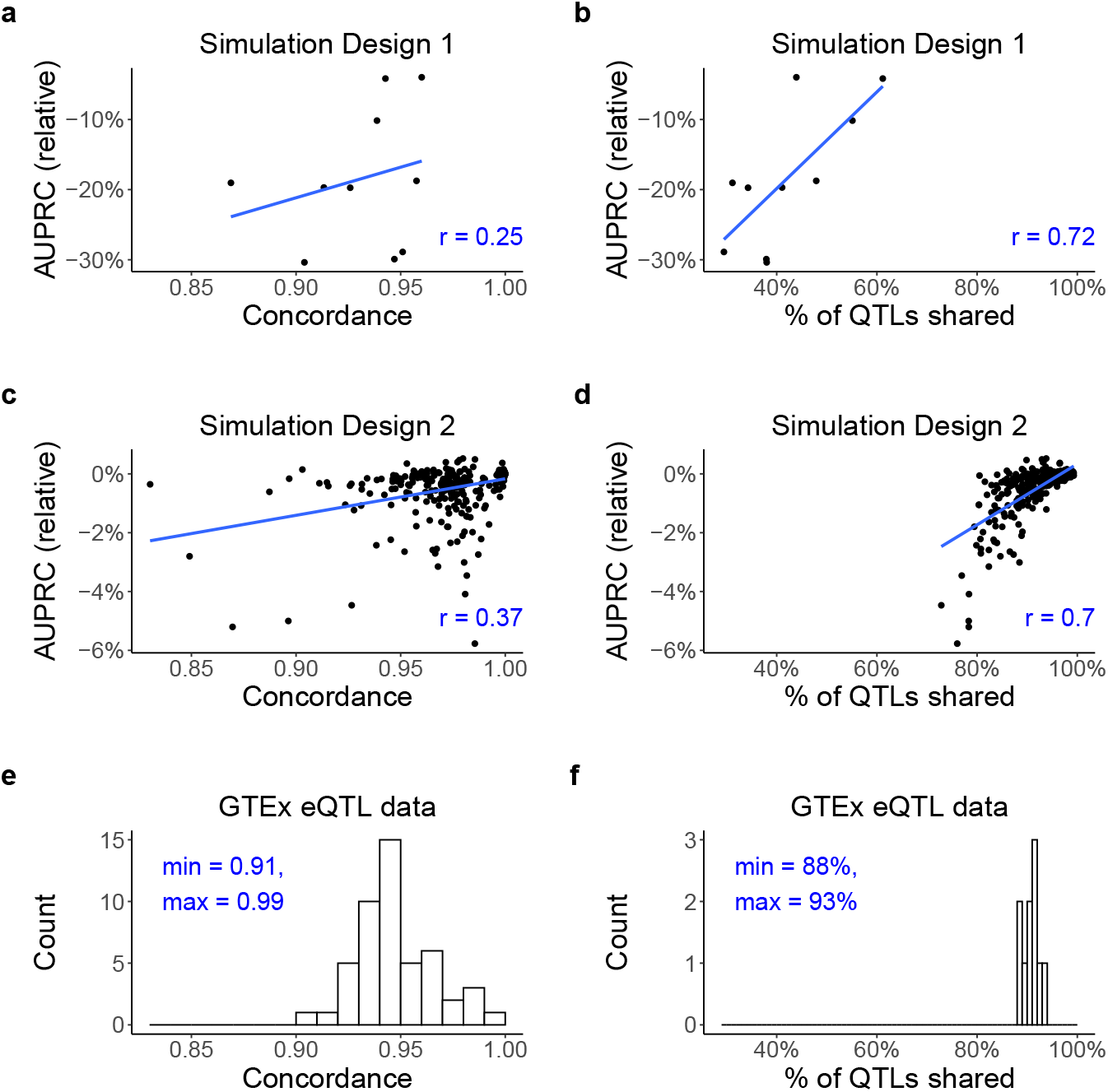
Joint analysis of results from Simulation Design 1, Simulation Design 2, and GTEx eQTL data [10]. The *x*-axes of **a, c**, and **e** show the concordance between PEER factors and top PCs (defined analogously as the concordance score; Section 4.1). The *x*-axes of **b, d**, and **f** show the percentage of QTL discoveries shared between PEER and PCA (in **b** and **d**, for each method, binary decisions are made based on *p*-values using the Benjamini-Hochberg (BH) procedure and a target false discovery rate of 0.05). In **a** through **d**, the *y*-axes show (AUPRC_PEER_ −AUPRC_PCA_) */*AUPRC_PCA_, the blue lines are the simple linear regression lines, and the Pearson correlation coefficients are shown on the bottom right. **a** and **b** each contains 10 data points, corresponding to the 10 simulated data sets in Simulation Design 1. **c** and **d** each contains 352 data points, corresponding to the 352 simulated data sets in Simulation Design 2. The methods compared in **a** through **d** are PCA direct screeK and PEER noCov trueK factors. **e** presents similar information as Figure 5; the total count is 49, which is the number of tissue types with GTEx eQTL analyses. **f** is based on Figure S6(d); the total count is 10, which is the number of tissue types randomly selected for analysis in Figure S6. We find that the percentage of QTL discoveries shared is a good predictor of the relative performance of PEER versus PCA and is a better predictor than concordance. This plot is also evidence that Simulation Design 2 is more realistic than Simulation Design 1 because the ranges that concordance and percentage of QTL discoveries shared fall in in **e** and **f** agree better with those in **c** and **d** than those in **a** and **b**.

### S5 Theory of PCA and HCP

#### S5.1 Principal component analysis (PCA)

Principal component analysis (PCA) [34, 35] is a well-established dimension reduction method with many applications. Here we aim to provide a brief summary of its algorithm, derivation, and interpretation.

Let *X* denote the *n* × *p* observed data matrix that is observation by feature, e.g., a molecular phenotype matrix. We use *X* instead of *Y* here to be consistent with standard PCA notations. We assume that the columns of *X* have been centered and scaled. That is, *X* satisfies

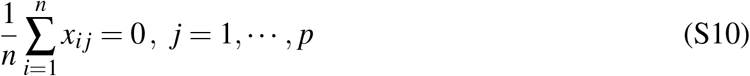

and

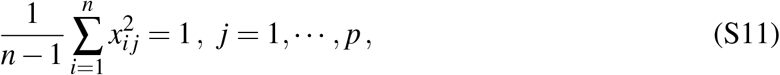

where *x*_*i j*_ denotes the *i j*-th entry of *X*.

The PCA algorithm consists of two steps. In the first step, we calculate the sample covariance matrix 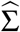 and perform eigendecomposition on it:

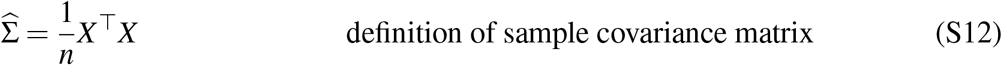

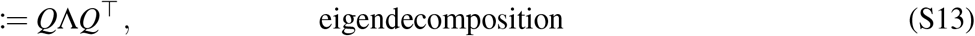

where

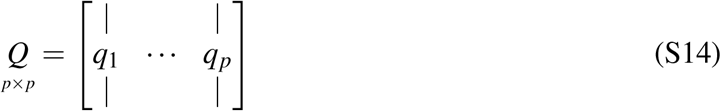

is an orthogonal matrix whose columns are eigenvectors of 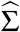, and

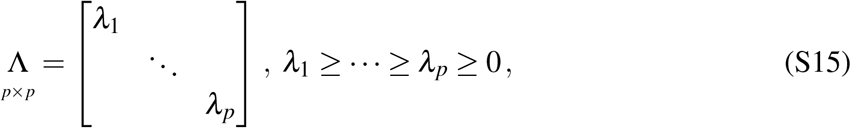

is a diagonal matrix whose diagonal entries are the corresponding eigenvalues of 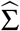. We know that 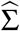 is orthogonally diagonalizable because it is a symmetric matrix (recall the spectral theorem for real matrices [55]: a matrix is orthogonally diagonalizable if and only if it is symmetric). The eigenvalues are all non-negative because 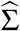 is positive semidefinite.

In the second step, we calculate *Z* as

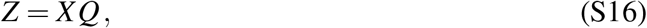

where the columns of *Z* are called the *principal components* (PCs) or *scores*, and *Q* is called the *loading matrix* or *rotation matrix*. It is worth noting that some authors may refer to *q*_1_, …, *q*_*p*_ as the PCs. This use of terminology is confusing and should be avoided [36].

The above two steps conclude the PCA algorithm. In practice, however, singular value decomposition (SVD) of the data matrix is often used as a more computationally efficient way of finding the loading matrix and the PCs [34].

The most common derivation of PCA is based on maximum variance [37]. First, we define 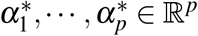 sequentially as

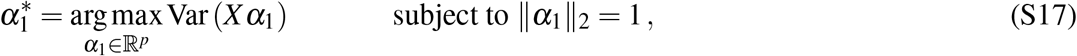

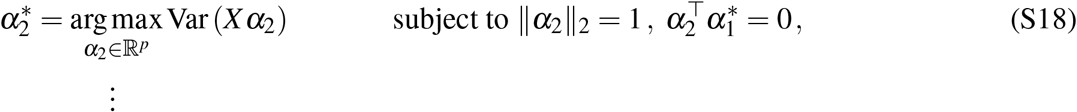

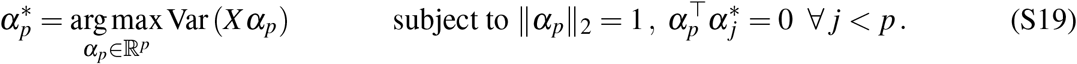

Then, we define the PCs of *X* as 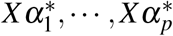. That is, the PCs are defined sequentially as the linear combinations of the columns of *X* with maximum variances, subject to certain constraints. It can then be shown that 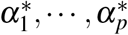 are given by *q*_1_, …, *q*_*p*_ respectively, where *q*_1_, …, *q*_*p*_ are eigenvectors of 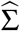 as defined in (S14).

A complementary property of PCA, which is closely related to the original discussion of Pearson [38], is the minimum reconstruction error property. Given *K < p*, we define *Q*_*K*_ as the matrix that contains the first *K* columns of *Q*. That is,

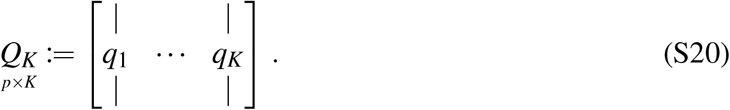

The minimum reconstruction error property of PCA states that *Q*_*K*_ is a global minimizer of the loss function

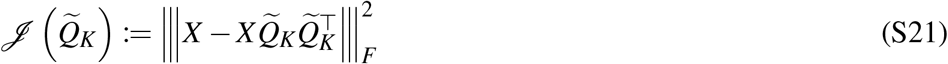

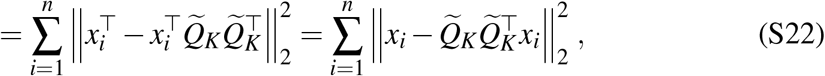

where 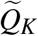 denotes an arbitrary *p* × *K* matrix whose columns are orthonormal, |||· |||_*F*_ denotes the Frobenius norm of a matrix, and 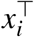 denotes the *i*th row of *X*. Since 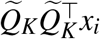 represents the (orthogonal) projection of *x*_*i*_ onto the subspace spanned by the columns of 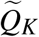, (S22) measures the total squared *ℓ*_2_ error when approximating each *x*_*i*_ with its projection onto the subspace spanned by the columns of 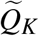.

A central idea of PCA is the proportion of variance explained by each PC. To establish this concept, we claim that

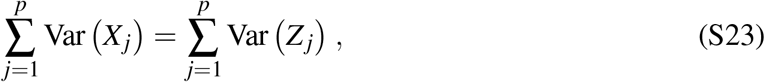

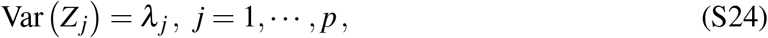

and

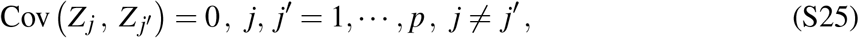

where *X*_*j*_ denotes the *j*th column of *X* (the *j*th original variable) and *Z*_*j*_ denotes the *j*th column of *Z* (the *j*th PC). (S25) means that the PCs are uncorrelated with each other.

We prove (S24) and (S25) by calculating 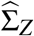, the sample covariance matrix of *Z* (we know that the columns of *Z* are centered by (S10) and (S16)):

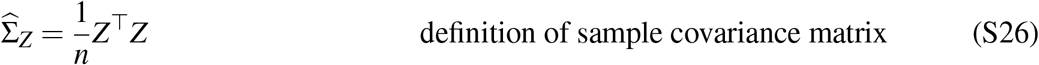

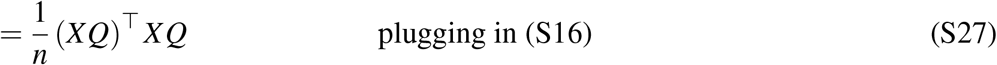

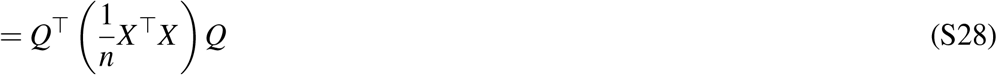

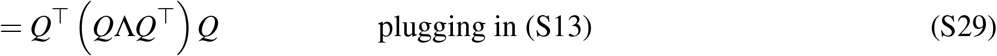

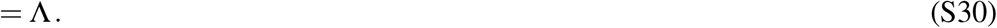

(S23) can be proven by the following:

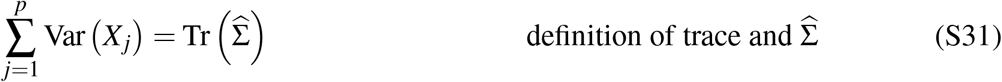

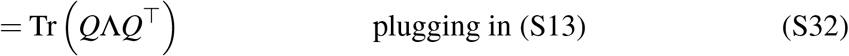

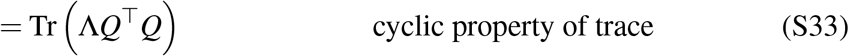

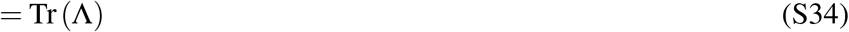

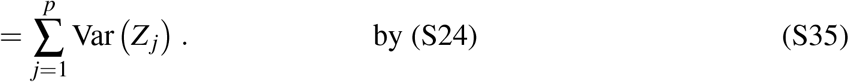

Because of (S23) and (S24), we may define the proportion of variance in the original data explained by the *j*th PC as

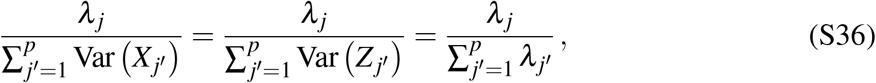

which provides a basis for deciding the number of PCs to keep (e.g., Algorithms S2 and S3).

#### S5.2 Hidden covariates with prior (HCP) and its connection to PCA

Hidden covariates with prior (HCP) [33] is a popular hidden variable inference method for QTL mapping defined by minimizing a loss function. Neither Mostafavi et al. [33] nor the HCP package documents the HCP method well. For example, the squares in the loss function (S37) are missing in both Mostafavi et al. [33] and the package documentation, but one can deduce that the squares are there by inspecting the coordinate descent steps in the source code of the R package. Here we aim to provide a better, more accurate documentation of the HCP method and point out its connection to PCA.

Given *Y*, the molecular phenotype matrix (*n* × *p*, sample by feature), *X*_1_, the known covariate matrix (*n* × *K*_1_, sample by covariate), *K*, the number of inferred covariates (HCPs), and *λ*_1_, *λ*_2_, *λ*_3_ *>* 0, the tuning parameters, HCP looks for

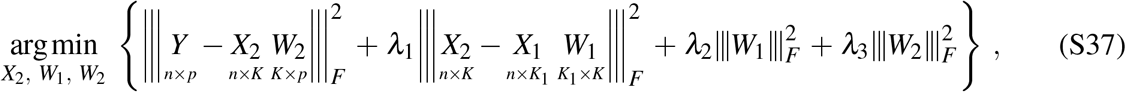

where |||· |||_*F*_ denotes the Frobenius norm of a matrix, *X*_2_ is the hidden covariate matrix, and *W*_1_ and *W*_2_ are weight matrices of the appropriate dimensions. The name of the method, “hidden covariates with prior”, comes from the second term in (S37), where the method informs the hidden covariates with the known covariates. The optimization is done through coordinate descent with one deterministic initialization (see source code of the HCP R package [33]). The columns of the obtained *X*_2_ are reported as the HCPs.

From (S37), we see that the HCP method is closely related to PCA. The first term in (S37) is very similar to (S21), the only difference being that the rows of *W*_2_ in (S37) are not required to be orthonormal and *X*_2_ is not required to be equal to 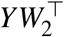.

##### Algorithm S2: The elbow method for choosing *K* in PCA (based on distance to diagonal line)

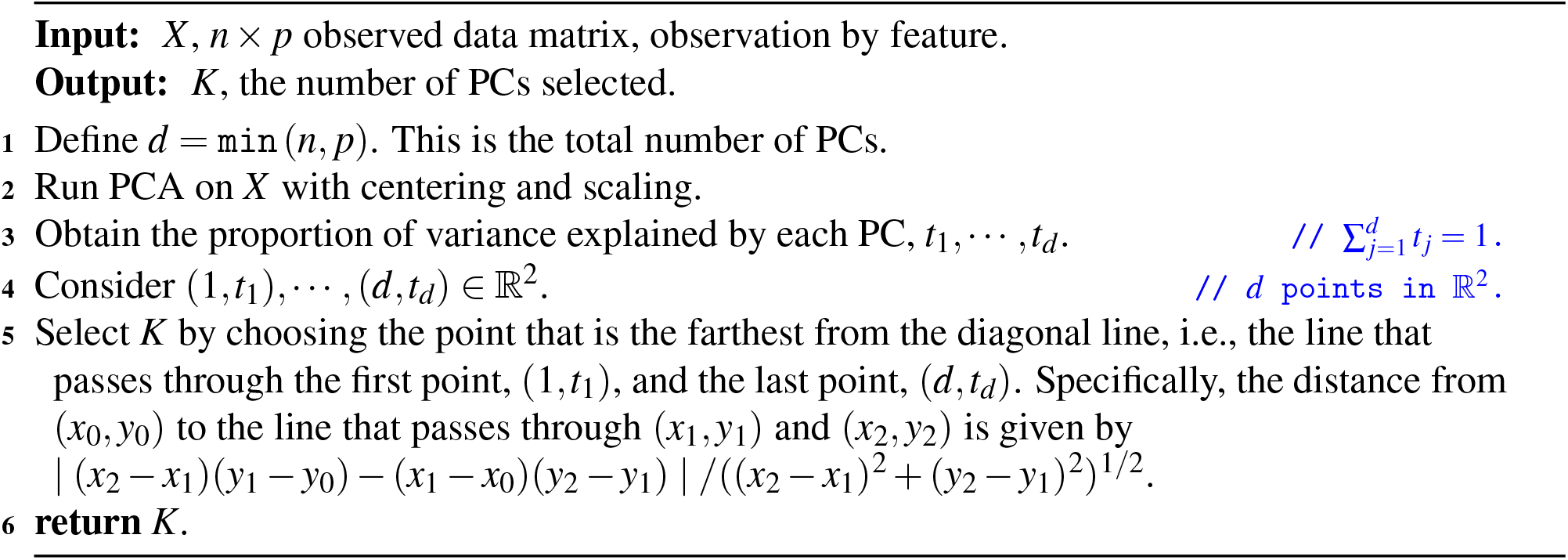

##### Algorithm S3: The Buja and Eyuboglu (BE) algorithm for choosing *K* in PCA

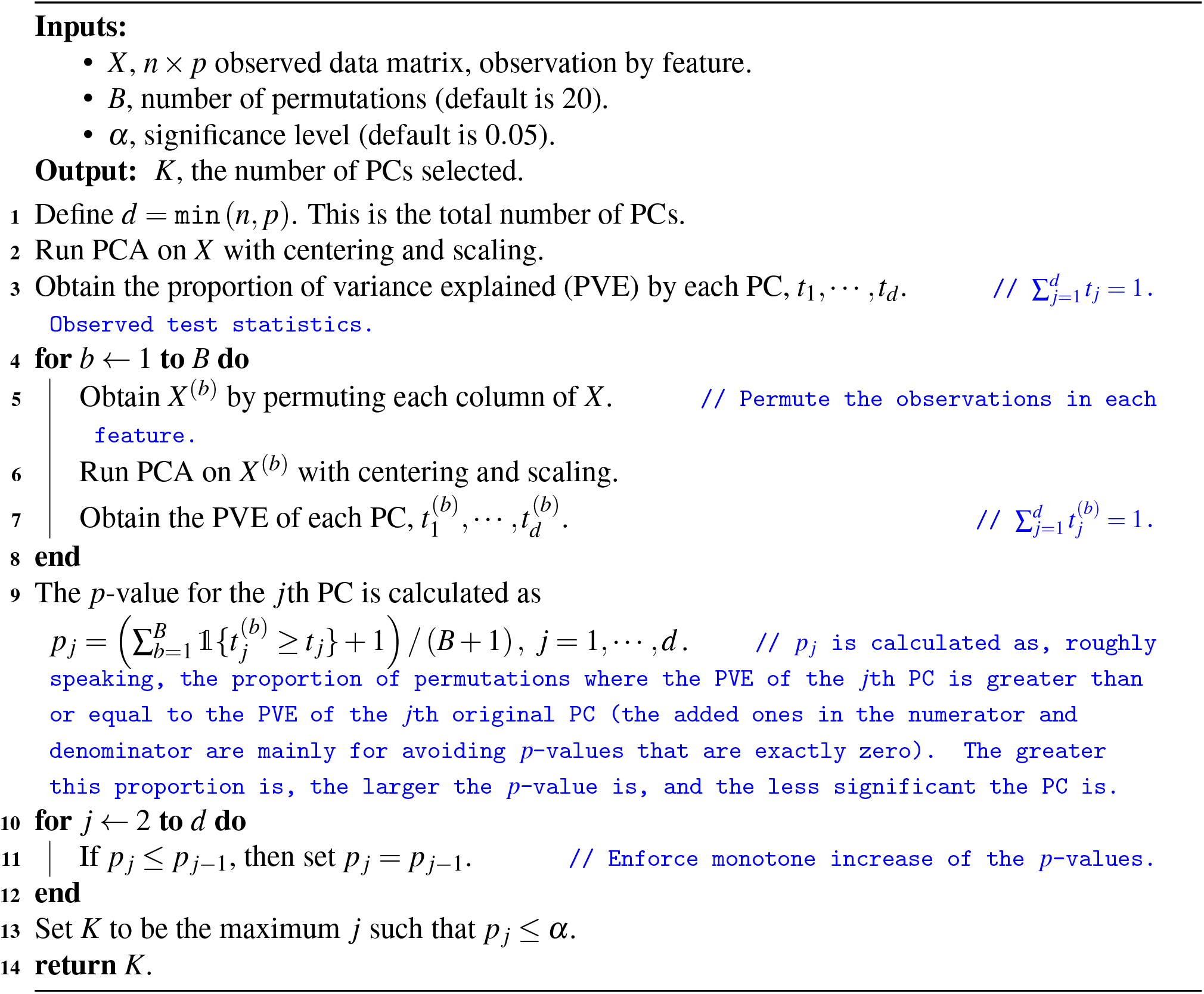

